# CAMSAP3 is required for mTORC1-dependent ependymal cell growth and lateral ventricle shaping in mouse brains

**DOI:** 10.1101/2020.07.19.211383

**Authors:** Toshiya Kimura, Hiroko Saito, Miwa Kawasaki, Masatoshi Takeichi

## Abstract

Microtubules (MTs) regulate numerous cellular processes, but their roles in brain morphogenesis are not well known. Here we show that CAMSAP3, a non-centrosomal microtubule regulator, is important for shaping the lateral ventricles. In differentiating ependymal cells, CAMSAP3 became concentrated at the apical domains, serving to generate MT networks at these sites. *Camsap3*-mutated mice showed abnormally narrow lateral ventricles, in which excessive stenosis or fusion was induced, leading to a decrease of neural stem cells at the ventricular and subventricular zones. This defect was ascribed at least in part to a failure of neocortical ependymal cells to broaden their apical domain, a process necessary for expanding the ventricular cavities. mTORC1 was required for ependymal cell growth but its activity was downregulated in mutant cells. Lysosomes, which mediate mTORC1 activation, tended to be reduced at the apical regions of the mutant cells, along with disorganized apical MT networks at the corresponding sites. These findings suggest that CAMSAP3 supports mTORC1 signaling required for ependymal cell growth via MT network regulation, and, in turn, shaping of the lateral ventricles.

**Summary statement:** CAMSAP3, which mediates non-centrosomal microtubule assembly, is required for mTORC1-dependent maturation of ependymal cells at the neocortex of developing mouse brains. Loss of CAMSAP3 causes deformation of the lateral ventricles.

## Introduction

Microtubules (MTs) are fundamental for a wide range of cellular processes such as cell division, polarization, migration, differentiation and growth. MT disruption dysregulates these processes, which leads to severe abnormalities in cell and/or body functions, often with fatal consequences. Clinically, MT-related alterations cause developmental disorders such as double cortex syndrome (Lasser et al., 2018). MTs are classified into two major categories, centrosomal and non-centrosomal. Non-centrosomal MTs emanate from sites other than the centrosome, such as the Golgi apparatus, nuclear envelope and apical cortex (Wu and Akhmanova, 2017). Although general functions of MTs in cellular and developmental processes have been well studied, specific roles of the non-centrosomal MT subsets are less known.

The CAMSAP/Nezha/Patronin family proteins are responsible for assembly of non-centrosomal MTs (Baines et al., 2009; Goodwin and Vale, 2010; Jiang et al., 2014; Meng et al., 2008; Tanaka et al., 2012). These proteins specifically bind to the MT minus ends, and are localized at various subcellular sites to which they tether MTs, resulting in formation of MT networks distinct from those organized by the centrosomes. For example, CAMSAP3 is localized at the apical cortex of intestinal epithelial cells, producing apicobasal arrays of MTs in these cells (Muroyama et al., 2018; Toya et al., 2016). CAMSAP2 and CAMSAP3 are differentially distributed in neuronal dendrites and axons, controlling their extension and polarity (Pongrakhananon et al., 2018; Yau et al., 2014). CAMSAP2 is also required for endothelial cell polarization in zebrafish (Martin et al., 2018), and Patronin is a key determinant of the anterior-posterior axis in *Drosophila* oocytes (Nashchekin et al., 2016). Thus, evidence has accumulated that this family of proteins plays many important roles in regulation of cell structure and function, but it remains unknown how much their functions contribute to the generation of higher-order structures such as tissues and organs.

The brain ventricles are cavities that are filled with cerebrospinal fluid (CSF). CSF circulates through multiple compartments of the ventricles, supporting the homeostasis of neurons and glia, and also maintaining quiescent adult neural stem cells (NSCs) (Delgado et al., 2014; Kokovay et al., 2012; Lowery and Sive, 2009). A major pool of adult NSCs are located in the zones adjacent to the surface of lateral ventricles, called the ventricular-subventricular zone (V-SVZ), where NSCs directly contact the CSF with their apical processes (Lim and Alvarez-Buylla, 2016; Mirzadeh et al., 2008). Once the ventricle is stenosed or closed, CSF circulation is occluded, leading to hydrocephalus (Kahle et al., 2016; Kousi and Katsanis, 2016; Lowery and Sive, 2009). At the areas of closure, neurogenesis is no longer maintained in the adult V-SVZ (Shook et al., 2012). The ventricles are lined with ependymal cells that develop multiple cilia on their apical surfaces, and these cilia play a crucial role in CSF circulation (Banizs et al., 2005; Ibañez-Tallon et al., 2004; Sawamoto et al., 2006).

In the present study, we show that CAMSAP3 is important for lateral ventricle shaping. Through phenotypic analysis of a mouse line in which *Camsap3* is mutated so as to abolish its function, we found that CAMSAP3 dysfunction causes various defects at the lateral ventricles, including enhancement of their closure and depletion of adult NSCs. Exploring mechanisms underlying these defects showed that CAMSAP3 supports mTORC1 signaling, which is important for the growth of ependymal cells, possibly through control of MT-dependent lysosomal positioning. Loss of this mechanism inhibited the normal expansion of the lateral ventricles, leading to their increased closure and associated events. These observations suggest that the CAMSAP3–non-centrosomal MT system is indispensable for constructing robust lateral ventricles, which support brain homeostasis.

## Results

### Increased ventricle stenosis and fusion in CAMSAP3 mutant brains

We used a CAMSAP3 mutant mouse, *Camsap3^dc/dc^*, in which the coding sequence for the C-terminal CKK domain of CAMSAP3, required for its binding to MTs, had been deleted from both alleles (Toya et al., 2016). Brains from adult mutant mice were slightly smaller than those from wild-type (WT) mice, having distinctly smaller olfactory bulbs (Figures 1A-C). Histological analysis of these brains at postnatal day (P) 28.5 showed no obvious gross defects in their cytoarchitecture, except for ventricular abnormalities (Figures 1D-F). In WT brains, the lateral ventricle was widely open at the dorsal regions, with narrowing at more ventral regions where stenosis or fusion occurred to variable extents among the individuals. The lateral ventricles in mutant brains exhibited much smaller cavities at the dorsal regions and more extensive stenosis or fusion at the ventral regions than in WT brains. Such ventricular defects were not clearly detectable in fetal brains (Figure S1), suggesting that the observed ventricular defects were elicited at postnatal stages. Immunostaining WT brains at P28.5 for CAMSAP3 showed that this protein is concentrated along the ependymal cell zones that line the lateral ventricles, and its mutated counterpart showed a similar distribution (Figure 1G), which implied that the CAMSAP3 mutant phenotypes observed above might be related to ependymal cell structure or function.

**Figure 1.**
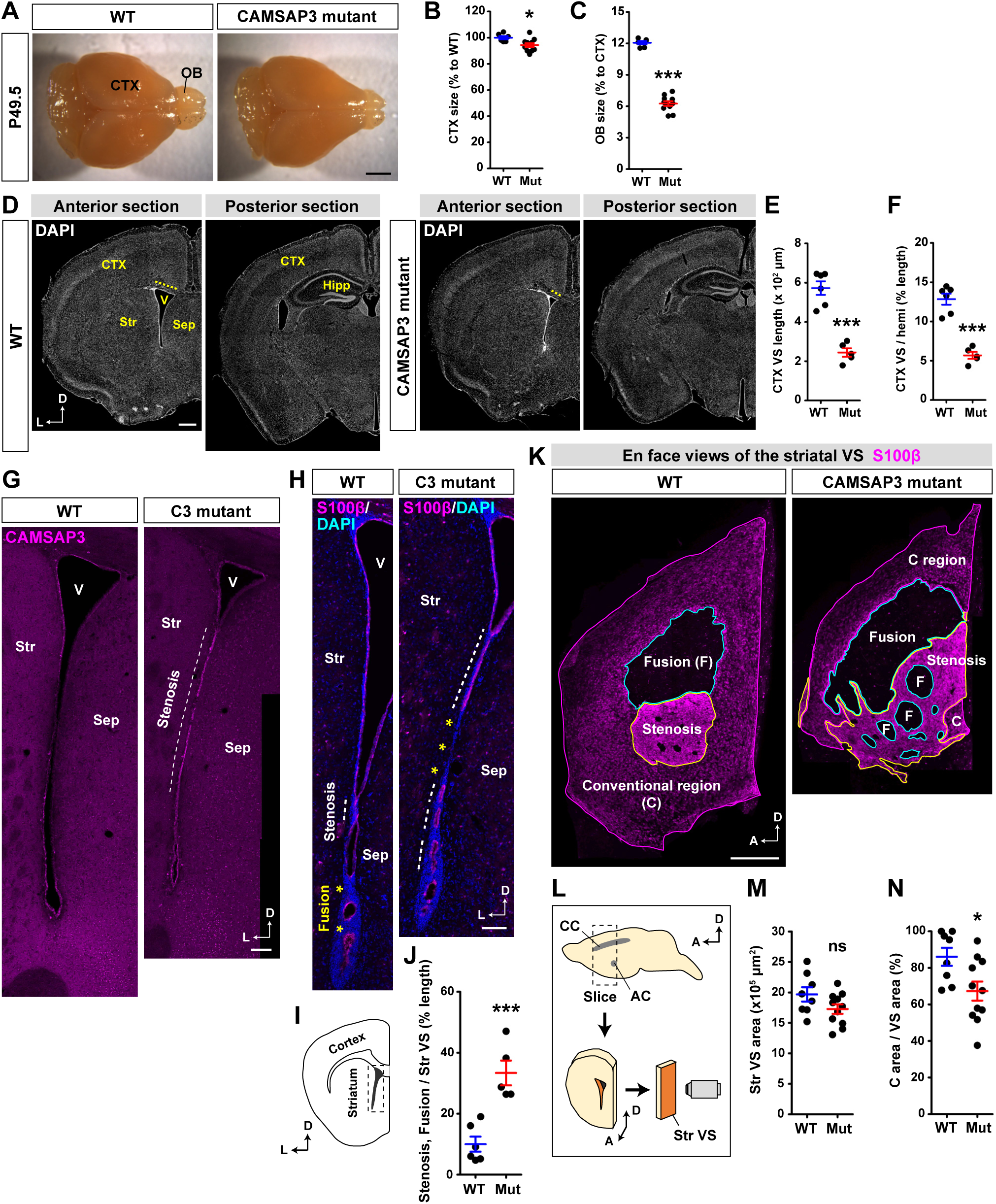
Accelerated ventricle stenosis and fusion in CAMSAP3 mutant brains. (**A**) Dorsal views of the whole forebrain at P49.5. Note the smaller olfactory bulb (OB) in the CAMSAP3 mutant brain. (**B**, **C**) Relative size of WT and mutant neocortices (B), and relative size of the OB to the neocortex (C) at P49.5. WT, n = 8; mutant, n = 12. (**D**) Coronal sections of the telencephalon at P28.5 stained with DAPI. Dotted lines indicate the mediolateral width of the neocortical VS. (**E, F**) Mediolateral width of the neocortical VS (E), and relative mediolateral width of the neocortical VS to the entire hemisphere (hemi) (F) at P28.5. WT, n = 6; mutant, n = 5. (**G**) Coronal sections of the telencephalon at P28.5, immunostained for CAMSAP3. Dotted line indicates stenosed regions. (**H**, **I**) The striatal VS in coronal sections of the telencephalon at P28.5 immunostained for S100β and with DAPI. The area of the striatal VS shown here is schematically explained in I. Dotted lines and asterisks indicate the stenosed and fused regions, respectively. (**J**) Proportion of the combined length of the stenosed and fused regions to the entire striatal VS length. WT, n = 6; mutant, n = 5. (**K, L**) En face view of striatal VS whole-mounts at P28.5 immunostained for S100β. Dorsal is up; anterior to the left. In L, schematic explanation of preparation of the specimens. An anterior part of the telencephalon was sliced at the position indicated by a box, and the slice containing the striatal VS (shown in orange) was further dissected. The VS was then viewed as drawn in the illustration. (**M**, **N**) Total area of the conventional, stenosed and fused regions on the striatal whole-mounts (M), and proportion of the conventional area to the total VS area. WT, n = 8; mutant, n = 11. L, lateral; D, dorsal; A, anterior; CTX, neocortex; OB, olfactory bulb; V, ventricle; Str, striatum; Sep, septum; Hipp, hippocampus; CC, callosal commissure; AC, anterior commissure; F, fusion. Error bars indicate SEM. ns, not significant, *p < 0.05, ***p < 0.001, Student’s t-test. Scale bars, 2 mm in (A), 500 μm in (D), 100 μm in (G) and (H), and 300 μm in (K). See also Figures S1 and S2.

Therefore, we closely examined the layers of ependymal cells by immunostaining for an ependymal cell marker, S100β (a calcium binding protein). In WT ventricular surfaces (VSs), S100β was detected through the VSs. In mutant brains, however, the regions lacking S100β^>^ ependymal cells were increased (Figures 1H-J). We also used another marker, parvalbumin (PV), which is specifically expressed in ependymal cells at the stenosed region (Filice et al., 2017), and found that PV-positive cells were increased in mutant brains (Figures S2A and S2B), confirming the enhanced stenosis in mutant brains. These results indicate that CAMSAP3 mutation promotes ventricular stenosis or fusion at the boundaries between the striatum and septum.

We also analyzed the anterior lateral ventricles using ‘en face’ views of the whole-mount striatum, identifying three distinct regions on the VS of both genotypes: #1) the S100β-negative region, #2) the region with strong, uniform S100β signals, and #3) the region with dotted S100β signals at moderate intensities (Figures 1K and 1L). Comparing these observations with those obtained using coronal sections, we can assume that region #1 corresponds to the fused region, region #2 to the stenosed region due to its PV expression (Figures S2C-E), and #3 to the ‘conventional’ ependymal cell layer that faces to the open ventricle. Analysis of these specimens showed that the fused and stenosed regions covered larger parts of the VSs in the mutant striatum than in WT, which resulted in a decrease of the conventional area in the mutants (Figures 1M and 1N). Ventricle stenosis and fusion frequently cause hydrocephalus (Kahle et al., 2016; Kousi and Katsanis, 2016), which is anatomically characterized by abnormal enlargement of the ventricle. However, CAMSAP3 mutant mice did not show any sign of ventricle enlargement at least in the telencephalon, rather they showed ventricle narrowing.

### Decrease of adult NSCs in the V-SVZ of the CAMSAP3 mutant striatum

Ependymal cells are part of the neural stem cell (NSC) niche at the ventricular and subventricular zone (V-SVZ) (Figure 2A) (Kokovay et al., 2008; Lim and Alvarez-Buylla, 2016; Paez-Gonzalez et al., 2011). Therefore, loss of ependymal cells could affect maintenance of adult NSCs. To determine whether adult NSCs are normally present at the striatal V-SVZ of CAMSAP3 mutant mice, we immunostained for VCAM1, a cell surface sialglycoprotein that had been proposed as a marker for adult NSCs (Kokovay et al., 2012). En face views (top views) of the striatal VSs in a whole-mount specimen confirmed that VCAM1 was exclusively localized on the surface of clustered cells having small apical domains (Figure 2B), which can be anatomically identified as NSCs. Immunostained coronal sections showed that VCAM1 signals were detected as puncta distributed along the VS in both WT and CAMSAP3 mutant brains at P28.5 (Figure 2C). These puncta closely associated with a tip of GFAP^+^ processes, which ensures that VCAM1 immunosignals represent parts of NSCs (Figure 2C, insert). Importantly, such VCAM1 signals were greatly reduced at the regions where stenosis or fusion was morphologically recognized, regardless of genotype, suggesting that NSCs were eliminated as a result of ventricle closure. Whole-mount samples confirmed that VCAM1^+^ clusters were mostly restricted to the conventional layer in both genotypes (Figures 2D and 2E). Additionally, even at the conventional area, the number of VCAM1^+^ clusters/area was decreased in about a half of the mutant mice examined (Figure 2F). Given that stenosis areas increased in mutant ventricles, we can estimate that the overall number of VCAM1^+^ clusters decreased in the mutants (Figure 2G). In addition, we noted that VCAM1^+^ cells had smaller apical domains in the mutants, although they formed clusters as WT cells did, and the number of cells per cluster did not change (Figures S3A-D), suggesting the possibility that CAMSAP dysfunction may also have affected morphology of NSCs.

**Figure 2.**
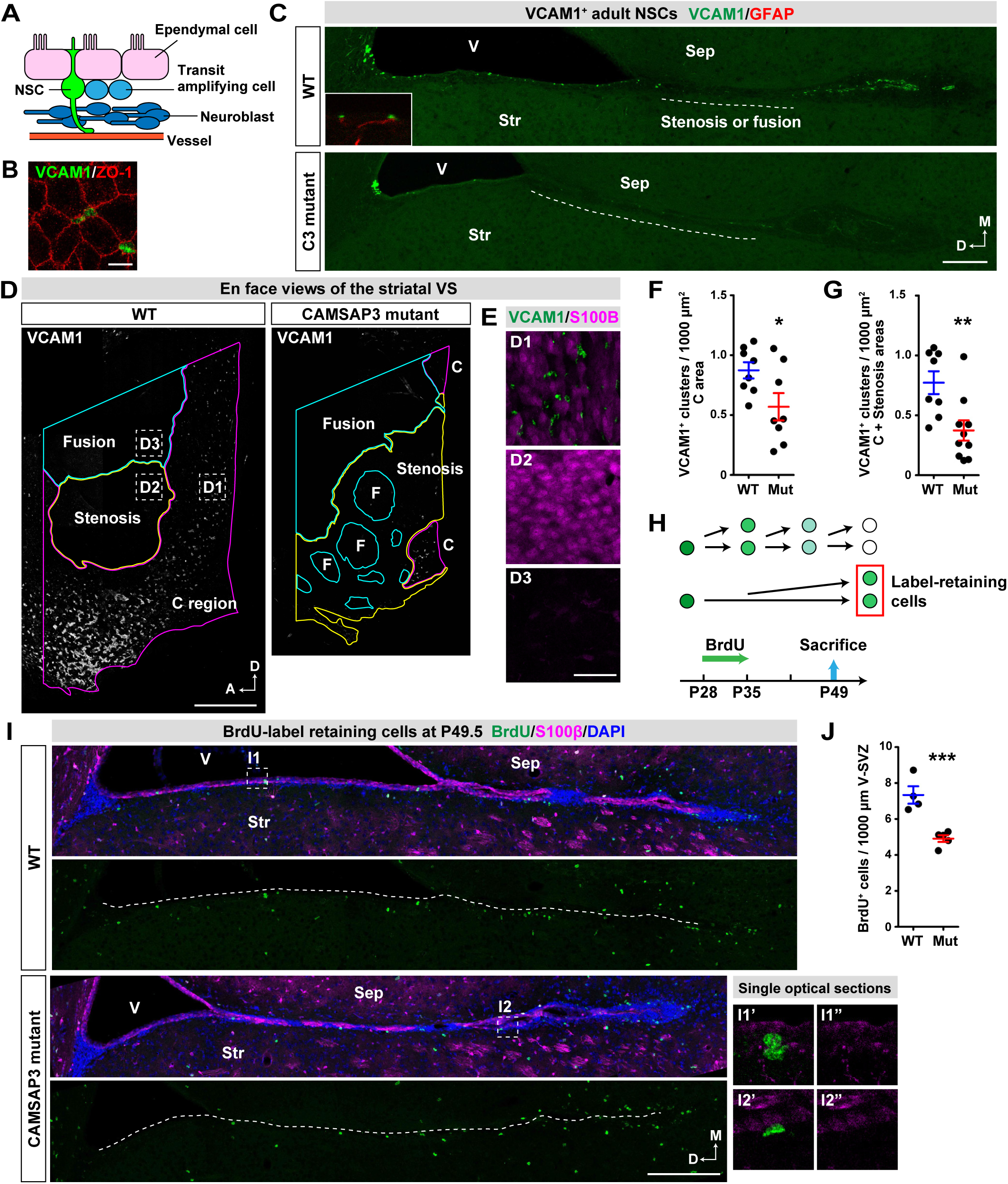
Decrease of adult NSCs at the striatal niche in CAMSAP3 mutant brains. (A) Schematic drawing of the adult NSC niche at the V-SVZ. NSCs are embedded in the ependymal cell layer and generate transit amplifying cells, which subsequently produce neuroblasts that migrate to the OB. The ventricle is up. (B) Top (en face) view of a striatal VS whole-mount at P28.5 immunostained for VCAM1 and ZO-1. (C) The striatal VS in coronal sections of the telencephalon at P28.5 immunostained for VCAM1. Dotted lines indicate the stenosed or fused regions. The inset shows the association of VCAM1^+^ puncta with GFAP^+^ processes. Medial is up, and dorsal to the left. (**D, E**) En face view of striatal VS whole-mounts at P28.5 immunostained for VCAM1. The same whole-mount samples as shown in Figure 1K were co-immunostained for VCAM1 and S100β. In E, the boxed regions are magnified to view VCAM1 and S100β at the conventional (D1), stenosed (D2) or fused (D3) regions. (**F**, **G**) The number of VCAM1^+^ clusters in the conventional region only (F), and in the conventional and stenosed regions (G). WT, n = 8; mutants, n = 8 (F), 10 (G). (H) Temporal difference in the BrdU label dilution due to cell divisions and the experimental timeline for BrdU administration. (I) Striatal V-SVZ regions in a coronal section of the telencephalon in P49.5 mice that were administered with BrdU, and co-stained for BrdU and S100β along with DAPI. Dotted lines indicate the striatal VS. The boxed regions are magnified at the lower right. (J) The number of BrdU^+^/S100β^-^ cells in the striatal V-SVZ. WT, n = 4; mutant, n = 5. M, medial; D, dorsal; A, anterior; V, ventricle; Str, striatum; Sep, septum; C, conventional region; F, fusion. Error bars indicate SEM. *p < 0.05, **p < 0.01, ***P = 0.0014. Student’s t-test. Scale bars, 10 μm in (B), 100 μm in (C), 300 μm in (D), 50 μm in (E) and 200 μm in (I). See also Figures S3 and S4.

We next examined adult NSCs using another approach, a BrdU-label retention assay. This assay exploits the slowly dividing nature of adult NSCs (Figure 2H) (Doetsch et al., 1999; Furutachi et al., 2015); once labeled, they retain the labels for a long period after the labeling protocol has ended. By contrast, rapidly dividing cells such as transit amplifying cells dilute the BrdU labels via multiple rounds of cell division. We administered P28.5 mice with BrdU for one week, followed by a non-administrating period for two weeks until sacrifice (Figure 2H). The results showed that fewer BrdU-retaining cells were detected in the V-SVZ of CAMSAP3 mutant striatum at P49.5 (Figures 2I and 2J), consistent with the idea that NSCs were decreased.

Adult NSCs in the V-SVZ produce interneurons that migrate to the olfactory bulb (OB) (Lim and Alvarez-Buylla, 2016). Therefore, we expected less production of OB interneurons in CAMSAP3 mutant mice, and this might explain why they have smaller OBs. To test this idea, we estimated the number of BrdU-retaining neurons in the entire granule cell layer (GCL) of the OB and found that the number was decreased in mutant OBs (Figure S4). Although the ratio of BrdU-retaining neurons to GCL neurons and the density of cells expressing NeuN, a neuronal marker, in the GCL were comparable between WT and mutant OBs (Figures S4B, S4C and S4F), the total BrdU-retaining neurons was decreased in mutant OBs (Figure S4E), due to the reduction in their overall size (Figures S4D and 1A-C). Together, these results consistently suggest that the morphological defects in the lateral ventricles resulted in a decrease of adult NSCs at the V-SVZ of CAMSAP3 mutant striatum.

### Failure of ependymal cells to grow normally at CAMSAP3 mutant neocortices

Then, we began to explore mechanisms by which loss of functional CAMSAP3 led to the narrowing of the lateral ventricle. This morphological change included shortening of the neocortical VSs along the mediolateral axis (Figure 1D). We paid attention to this abnormality, because we can presume that the reduced growth of the neocortical VSs would lead to narrower opening of the ventricular cavity. To seek the cellular basis of the shortening of neocortical VSs in the mutants, we first observed morphology of individual ependymal cells in P28.5 brains, by immunostaining for N-cadherin as a cell junction marker, as well as for γ-tubulin, a component of centrosomes or basal bodies (BBs), which are multiplied to produce multiple cilia in differentiated ependymal cells. We found that differentiation of ependymal cells normally took place at mutant neocortices, as assessed by appearance of a multiple BB cluster in each cell. Remarkably, however, their apical domain size was significantly reduced in the mutants (Figures 3A and 3B). Moreover, their number was also decreased in these brains (Figures 3C and 3D). Overall, the mediolateral length of VSs was reduced by 60% in the mutant neocortex at P28.5.

**Figure 3.**
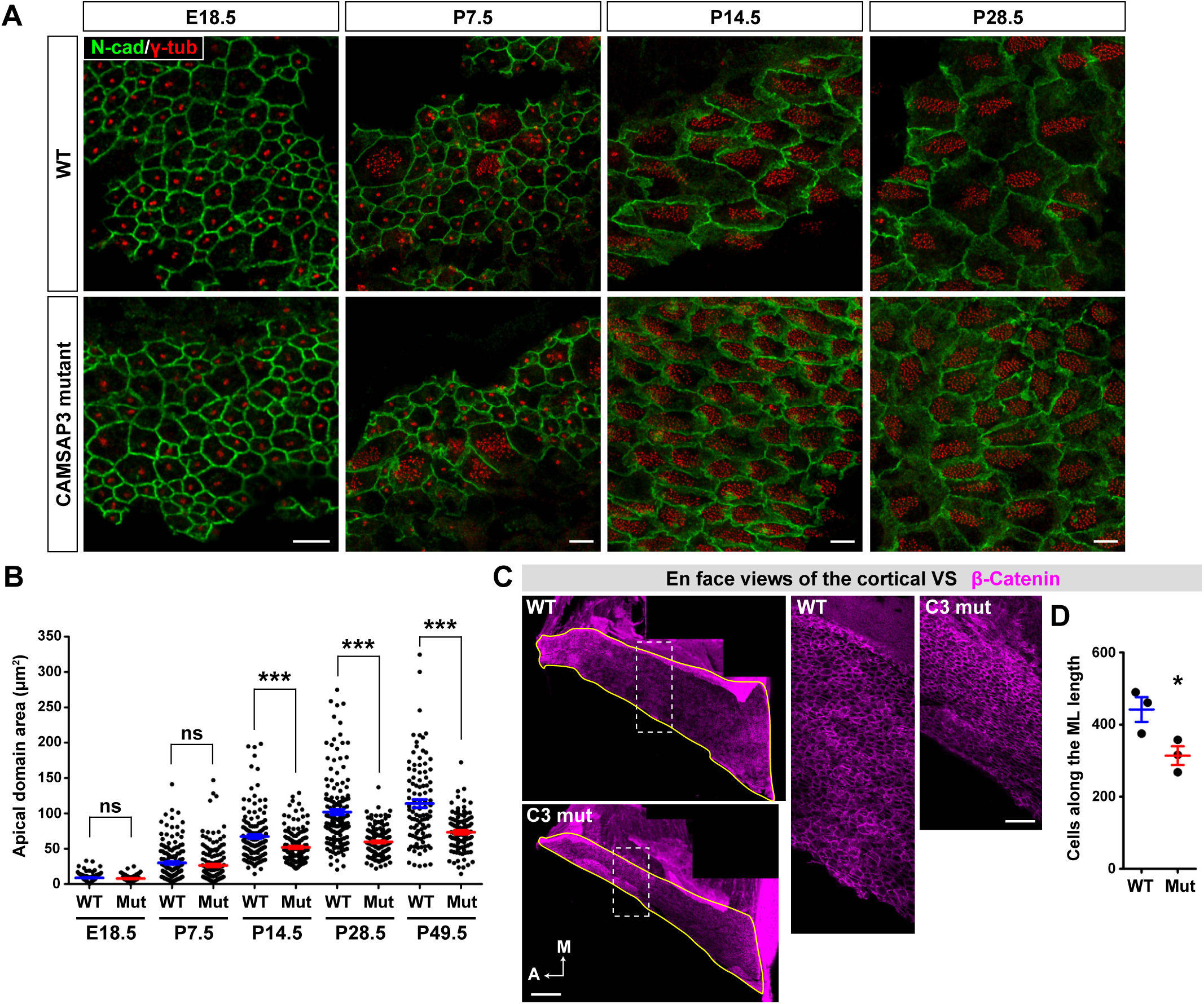
Decreases in the apical domain area of ependymal cells and in their number in CAMSAP3 mutant neocortices. (A) Top (en face) views of cells composing the neocortical VS at different ages, immunostained for N-cadherin and γ-tubulin. (B) Apical domain area of cells at the neocortical VS. n = 50 cells per brain. n = 3 brains for each genotype at P7.5, P14.5 and P28.5; n = 2 brains for each genotype at E18.5 and P49.5. (C) En face views of a neocortical VS whole-mount, immunostained for β-catenin. The boxed regions are magnified at the right. (D) The number of cells along the mediolateral length of the neocortical VS. WT, n = 3; mutant, n = 3. M, medial; A, anterior. Error bars indicate SEM. ns, not significant, *p < 0.05, ***p < 0.001, Student’s t-test. Scale bars, 5 μm in (A), 200 μm in the left panels of (C), and 50 μm in the right panels of (C). See also Figures S3.

The ependymal cell apical domain is known to enlarge during the postnatal period at the striatum in a planar cell polarity fashion (Redmond et al., 2019). We therefore examined whether this would also be the case at the neocortex by observing developmental processes of ependymal cells. At E18.5, a stage prior to ependymal cell differentiation, the neocortical VSs were solely covered by mono-ciliated radial glial cells, defined as ependymal precursor cells (Spassky et al., 2005) (Figure 3A). The apical domain size in these cells did not differ between WT and mutant brains (Figure 3B). At P7.5, centriole amplification had begun in some cells to produce multiple cilia, which was accompanied by apical domain growth in both genotypes (Figure 3A). Usually, the apical domain size was the smallest in cells with a monomeric BB, and it increased along with centriole amplification, reaching the maximum in cells that have acquired multiple BBs. The ratio of such differentiating cells (with multiple BBs) to undifferentiated cells (with a monomeric BB) did not greatly differ between the two genotypes, nor did their apical domain sizes differ at this stage (Figure 3B). At P14.5, however, a difference in the apical domain size between WT and mutant cells became evident. While the VSs at this age were covered primarily by multiciliated cells in both genotypes, those in mutant neocortices displayed narrower apical domain areas than WT cells (Figures 3A and 3B). By P28.5, apical domain growth was largely completed, and the difference in apical domain size between WT and mutant cells was thus fixed (Figures 3A and 3B). To summarize, apical domain growth in CAMSAP3-mutated ependymal cells was impaired during postnatal periods. Closer cytological observations further indicated that the number of BBs and the areas occupied by BBs at the apical domain of ependymal cells did not particularly differ between WT and mutant neocortices; that is, only the non-ciliated areas failed to grow normally in the mutants.

We also analyzed apical domain size in ependymal cells lining the striatum of P28.5 brains, at the dorsal portions which are not closed, and found that the size was only slightly reduced in the mutants (Figures S3E and S3F). We did not perform this analysis for the closed portions, as normal ependymal cells did not always persist there. Concerning the reduction in ependymal cell number in mutant neocortices, we could not determine how this occurred. Ependymal cell differentiation starts early in the postnatal period and, once differentiated, they become postmitotic (Spassky et al., 2005). We did not find any decrease in cell number at E18.5 and P7.5, suggesting that cell proliferation proceeded normally in the mutant neocortices. We also examined potential involvement of cell death. However, our immunostaining for cleaved Caspase-3 did not detect any clear apoptotic figures at neocortical ventricular zones in either WT or mutants at P7.5 and P28.5. We suspected that some delamination of ependymal cells might have been induced at the mutant neocortices, but we did not capture any images that support this idea.

### Downregulation of mTORC1 signaling in CAMSAP3 mutant ependymal cells

Previous work suggested that mTORC1 signaling regulates the apical domain size of radial glial cells (Foerster et al., 2017). We therefore examined whether an mTORC1-depedent mechanism is also involved in ependymal cell growth. We immunostained neocortical VSs of WT brains at various stages for phosphorylated S6 ribosomal protein (p-S6RP), an mTORC1 signaling activity readout (Figure 4A) (Ruvinsky and Meyuhas, 2006), observing its distribution. At E18.5, p-S6RP was only faintly detected in radial glial or ependymal precursor cells (Figure 4B). At P7.5, cytosolic p-S6RP became detectable in some of the ventricular cells (Figures 4B and 4C), and expression of active mTORC1 at this stage was also confirmed by Western blotting (Figure 4D). Triple immunostaining for p-S6RP, γ-tubulin and N-cadherin showed that p-S6RP was detected in enlarged cells, particularly in those having amplifying centrioles (Figures 4E-G), suggesting that activation of mTORC1 signals coincides with ependymal cell differentiation. In cells that had acquired multiple BBs, however, p-S6RP signals dropped, although some of them remained along the plasma membranes (Figures 4B and 4E-4G). These observations suggest that mTORC1 is transiently activated in developing ependymal cells at the WT neocortex. To test whether this activation of mTORC1 is important for apical domain growth of ependymal cells, we injected postnatal WT mice with the mTORC1 inhibitor rapamycin. Daily injections of the inhibitor from P6.5 for eight consecutive days resulted in a marked decrease of the apical domain size (Figure S5), confirming that mTORC1 signaling drives apical domain growth of ependymal cells.

**Figure 4.**
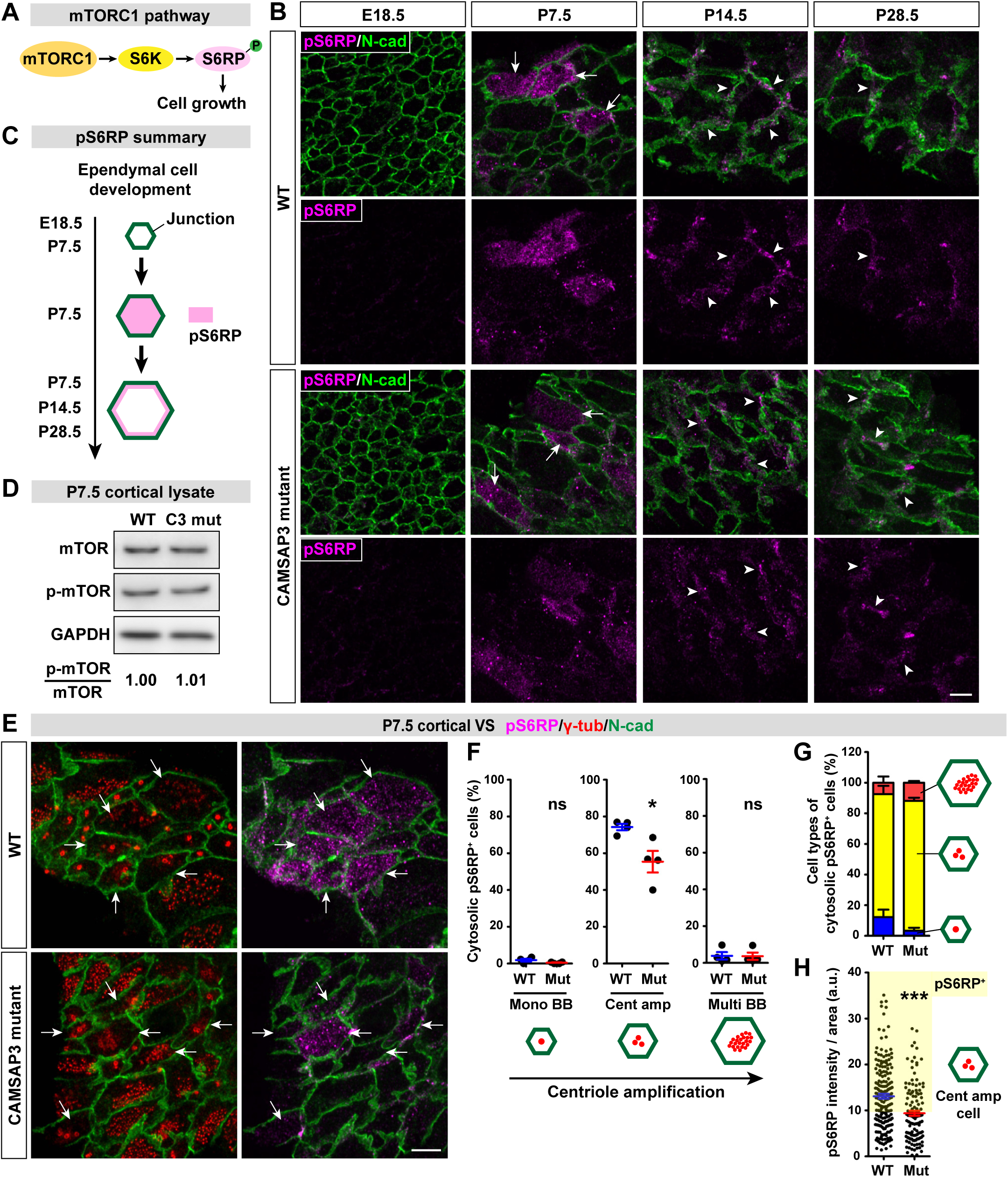
Downregulation of mTORC1 signals in developing ependymal cells of CAMSAP3 mutant neocortices. (A) mTORC1-S6RP signaling pathway. S6RP is activated by mTORC1 via S6 kinase (S6K) to promote cell growth. (B) Top views of cells at the neocortical VS at different ages, immunostained for phospho-S6RP and N-cadherin. Arrows and arrowheads point to cells with cytosolic and membrane-associated signals of phopho-S6RP, respectively. Phospho-S6RP signals were acquired using the same microscope setting and processed in the same way for all the panels. (C) Summary of phospho-S6RP distributions. S6RP phosphorylation is transiently elevated in the cytoplasm at P7.5, and thereafter held at the lateral plasma membranes. (D) mTOR phosphorylation detected by Western blotting of P7.5 neocortical lysates. Three brains were examined for each genotype and a representative image is presented. The values below the panels indicate the ratio of phospho-mTOR to the total mTOR. (E) Top views of cells at the neocortical VS at P7.5, immunostained for phospho-S6RP, γ-tubulin and N-cadherin. Arrows point to cells undergoing centriole amplification. (F) Proportion of cytosolic phospho-S6RP^+^ cells in the cell group with monomeric BB, amplifying centrioles or multiple BBs at P7.5. n = 1,250 cells in 4 brains for WT; n = 1,013 cells in 4 brains for mutant. (G) Proportion of cytosolic phospho-S6RP^+^ cells belonging to each of the indicated cell group in the total cytosolic phospho-S6RP^+^ cells. n = 1,250 cells in 4 brains for WT; 1,013 cells in 4 brains for CAMSAP3 mutant. (H) Phospho-S6RP signal intensity in cells undergoing centriole amplification at P7.5. n = 185 cells in 4 brains for WT; n = 125 cells in 4 brains for CAMSAP3 mutant. Cells with a higher signal intensity than 10 luminance (a.u., highlighted) were regarded as phospho-S6RP^+^ cells in (F) and (G). Error bars indicate SEM. ns, not significant, *p < 0.05, ***p < 0.001, Student’s t-test. Scale bars, 5 μm. See also Figure S5.

We then asked what happened to mTORC1 signals in CAMSAP3 mutant ependymal cells. Total phosphorylation level of mTOR was not much changed in the mutant neocortex (Figure 4D). Immunostaining of neocortical VSs for p-S6RP in mutant cells showed its distribution pattern was similar to that in WT cells in all the stages examined (Figures 4B and 4E-4G). However, the overall intensity of cytosolic p-S6RP immunosignals was significantly decreased in mutant cells (Figures 4E, 4F and 4H). Collectively, these results suggest that mTORC1 activity is downregulated in CAMSAP3-mutated ependymal cells at postnatal periods, although its overall activity in neocortical lysates did not differ between WT and mutants (Figure 4D).

As an alternative mechanism that may reduce the apical domain size in ependymal cells, we considered the possibility that the apical domain had actively constricted, a phenomenon that occurs widely during epithelial morphogenesis (Takeichi, 2014). Such constriction is generally executed by activation of myosin II, which is associated with the adherens junction–F-actin complex (Takeichi, 2014). However, immunostaining for an activated (phosphorylated) form of myosin regulatory light chain 2 did not show any difference in its distribution at the apical domains or junctions between WT and mutant cells at least in P28.5 brains, suggesting that this possibility is unlikely.

### Reduction of lysosomes at apical regions of CAMSAP3 mutant ependymal cells

mTORC1 is activated on lysosomes (Betz and Hall, 2013; Sancak et al., 2010) (Figure 5A), and we confirmed that mTORC1 is closely associated with lysosomes in ependymal cells (Figure 5B). Previous work showed that lysosomal positioning is critical for mTORC1 activation. For example, forced displacement of lysosomes from the plasma membranes was sufficient to downregulate mTORC1 signaling (Korolchuk et al., 2011), as signaling modules upstream of mTORC1 (such as Akt) accumulate beneath the plasma membranes (Finlay and Cantrell, 2011; Gao et al., 2014). In ependymal cells at P7.5, an activated form of Akt (phospho-Akt) was detected as punctate immunosignals not only at the centriole amplification stage but also at the multiple BB stage in both WT and CAMSAP3 mutant neocortices (Figures 5C and 5D). By shifting focus planes, we confirmed that phospho-Akt was most abundant at around the level of adherens junctions, which are located at subapical regions of the cell (Figures 5E and 5F). Other signaling pathways upstream of mTORC1, such as ERK and AMPK, were rarely or only faintly detected at the corresponding regions of these cells in both genotypes at P7.5.

**Figure 5.**
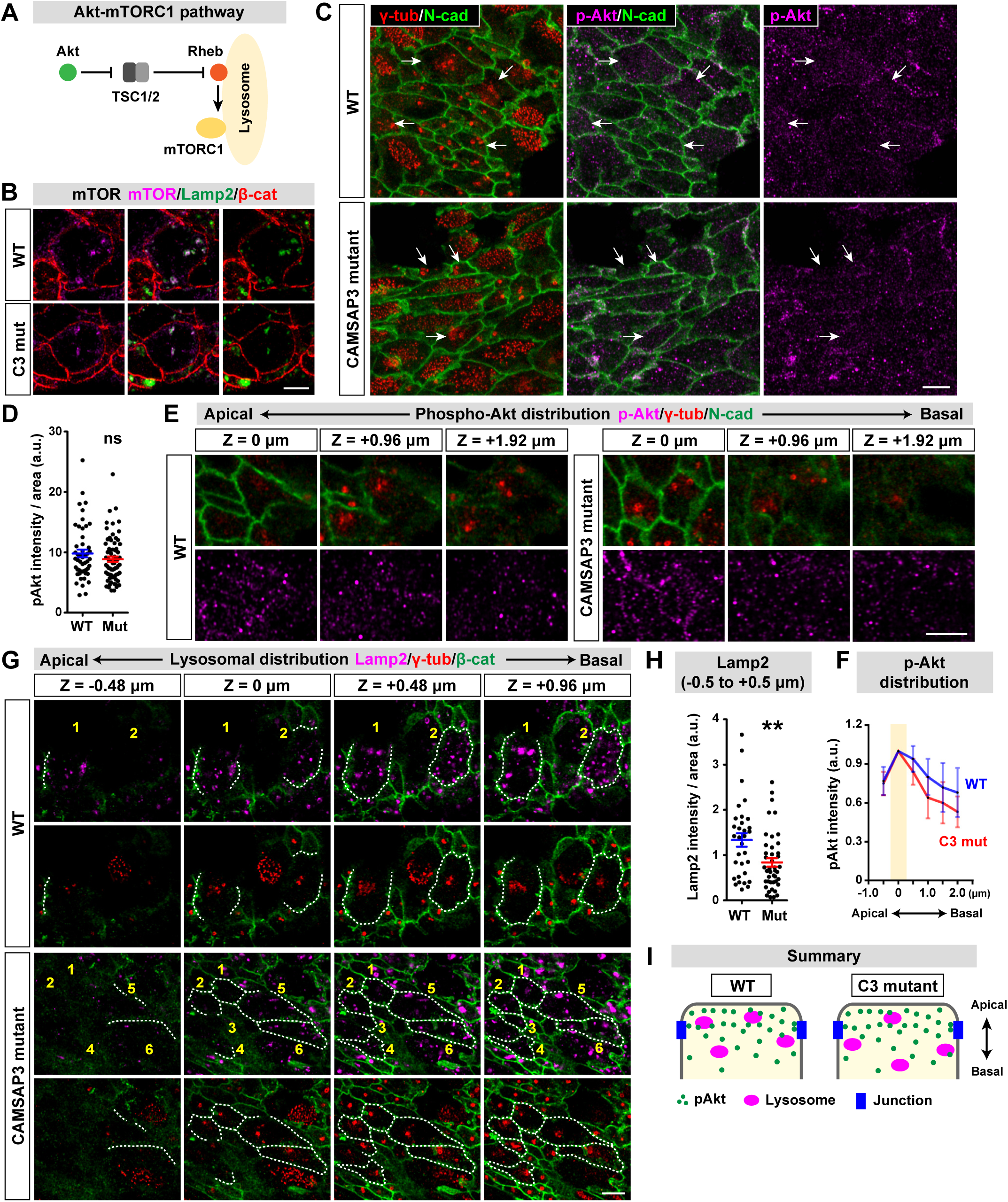
Distribution of phospho-Akt and lysosomes in developing ependymal cells of P7.5 neocortices. (A) Akt-mTORC1 signaling pathway. Akt liberates Rheb through inhibition of TSC1/2, resulting in activation of mTORC1 on lysosomal membranes. (B) Top views of cells at the neocortical VS, immunostained for mTOR, Lamp2 and β-catenin. (C) Top views of cells at the neocortical VS, immunostained for phospho-Akt, γ-tubulin and N-cadherin. Arrows point to cells undergoing centriole amplification. (D) Phospho-Akt immunostaining intensity in cells undergoing centriole amplification. n = 50 cells in 3 brains for WT; 70 cells in 3 brains for CAMSAP3 mutant. (E) Apicobasal distribution of phospho-Akt in cells undergoing centriole amplification. The neocortical VSs were immunostained for phospho-Akt, γ-tubulin and N-cadherin and viewed from the top. Single optical sections of 0.16 μm in thickness taken at different apicobasal levels are shown. (F) Analysis of the apicobasal distribution of phospho-Akt immunostaining signals in cells undergoing centriole amplification. The level on which the N-cadherin^+^ adherens junctions can be focused was set as 0 μm and highlighted. The staining intensity at 0 μm was normalized to 1.0. n = 35 cells in 3 brains for WT; 47 cells in 3 brains for CAMSAP3 mutant. (G) Apicobasal distribution of Lamp2^+^ lysosomes in cells undergoing centriole amplification. The neocortical VSs were immunostained for Lamp2, γ-tubulin and β-catenin and viewed from the top. Cells are numerically labeled. Single optical sections of 0.16 μm in thickness taken at different apicobasal levels are shown. Dotted lines trace β-catenin^+^ adherens junctions. (H) Lamp2 immunostaining intensity at the apical (−0.5 to +0.5 μm) domain of cells undergoing centriole amplification. The level on which the apical edges of β-catenin^+^ adherens junctions can be focused was set as 0 μm. Inclination of the VS to the optical plane was normalized (see Materials and Methods). n = 32 cells in 3 brains for WT; 44 cells in 3 brains for CAMSAP3 mutant. (I) Summary of the apicobasal distribution of phospho-Akt and Lamp2^+^ lysosomes in cells undergoing centriole amplification. Phospho-Akt is similarly enriched in the apical domain of the cells in both genotypes, whereas fewer lysosomes are present there in CAMSAP3 mutant cells. Error bars indicate SEM in (D) and (H), and SD in (F). ns, not significant, **p < 0.01, Student’s t-test. Scale bars, 5 μm.

We then examined the distribution of lysosomes, using the lysosomal marker Lamp2 in ependymal cells at P7.5, and found that Lamp2^+^ structures tended to decrease in the apical portions of mutant cells (Figures 5G-I). This may explain why mTORC1 activity was reduced in mutant ependymal cells.

### Depletion of apical MT networks in CAMSAP3 mutant ependymal cells

Lysosomes are transported along MTs (Bonifacino and Neefjes, 2017; Pu et al., 2016). Therefore, we investigated whether MT networks showed any changes due to CAMSAP3 dysfunction in neocortical ependymal cells. We first observed CAMSAP3 distribution at various developmental stages by immunostaining, finding that its staining signals were relatively weak at E18.5, but increased by P7.5, persisting to P28.5 (Figure S6A). Interestingly, the upregulation of CAMSAP3 selectively occurred in cells with either amplifying centrosomes or multiple BBs (Figure 6A and 6B), suggesting some functional relations between this upregulation and mTORC1 activation that also occurred during centriole amplification, although the upregulation of CAMSAP3 persisted to the multiple BB stage unlike mTORC1. CAMSAP3 was concentrated at the apical domain of a cell, more precisely, at a level more apical than N-cadherin^+^ adherens junctions (AJs) that are present at a subapical region of lateral cell membranes (Figures 6A and 6B). Comparison of CAMSAP3 and γ-tubulin localizations showed that, although γ-tubulin^+^ centrosomes or BBs were clustered at a particular region of the cytoplasm, CAMSAP3 was scattered throughout the apical cytoplasm, being located more apical than the cluster of basal bodies (Figures 6A and S6B). Mutated CAMSAP3 proteins were also concentrated at the apicalmost domain of cells with amplifying centrosomes or multiple BBs at P7.5 and later (Figures 6A and S6A).

**Figure 6.**
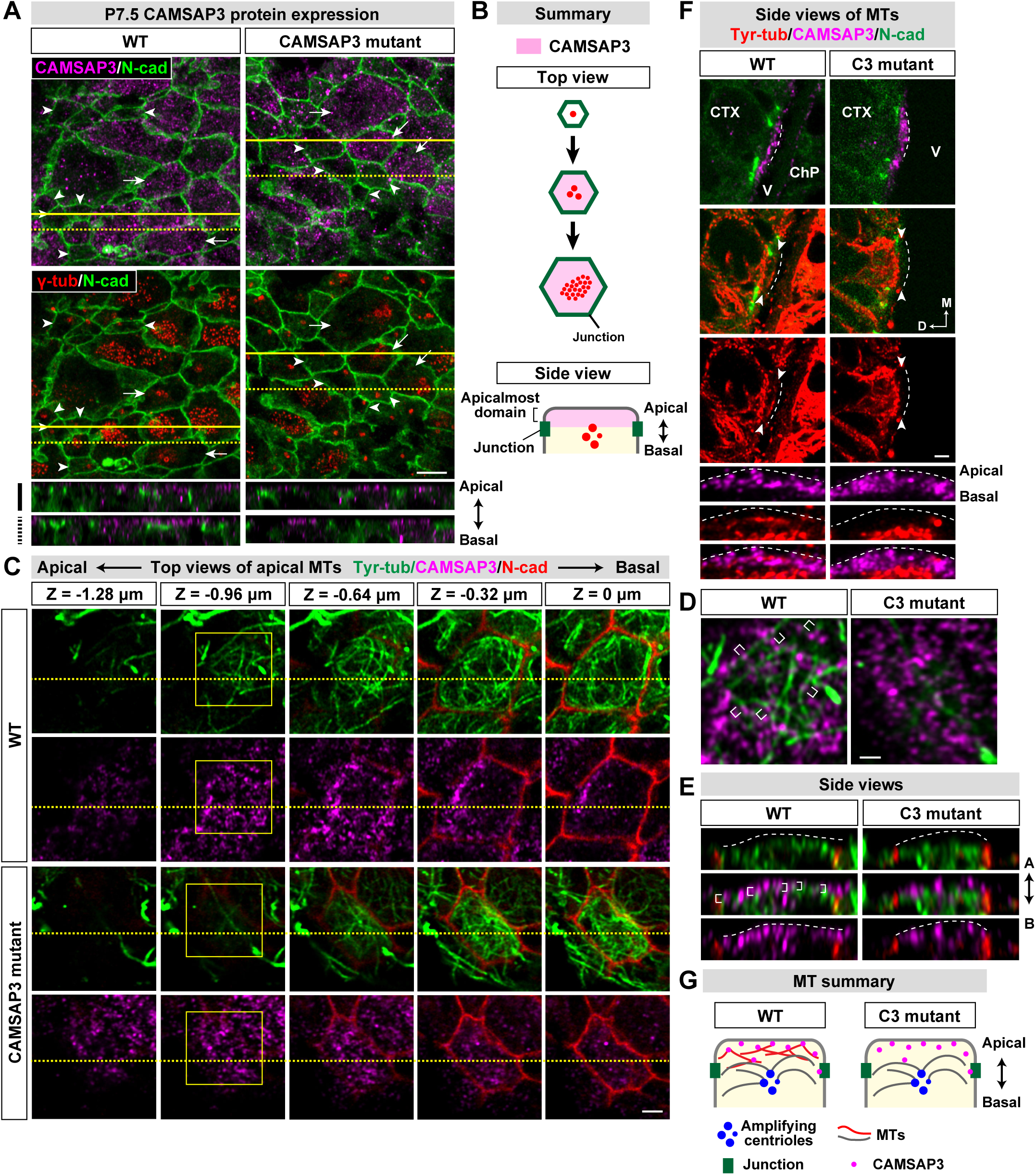
CAMSAP3 distribution and microtubule networks at the apicalmost regions of developing ependymal cells in P7.5 neocortices. (A) Top views of ependymal cells immunostained for CAMSAP3, γ-tubulin and N-cadherin. Arrows and arrowheads point to cells undergoing centriole amplification and those with the monomeric BB, respectively. Orthogonal (side) views at the solid and dotted lines are shown below. (B) Summary of CAMSAP3 distribution. Its expression is upregulated concurrently with centriole amplification and maintained thereafter. The protein is concentrated at the apicalmost domain, at a level more apical than N-cadherin^+^ adherens junctions. (C) Top views of ependymal cells immunostained for tyrosinated tubulin, CAMSAP3 and N-cadherin. Single optical sections of 0.16 μm in thickness taken at different apicobasal levels are shown. (D) Spatial relation between CAMSAP3 and MTs. The portions enclosed by yellow boxes in C are enlarged. Brackets point to representative colocalization of CAMSAP3 and MTs. (E) Orthogonal (side) views of ependymal cells at their apical regions. The images were collected along the yellow broken lines drawn in C. White dotted lines indicate the apical border of the CAMSAP3 puncta-containing zone. Brackets point to examples of the colocalization between CAMSAP3 and MTs. (F) Coronal (side) views of ependymal cells immunostained for tyrosinated tubulin, CAMSAP3 and N-cadherin. Cells whose apical surface was perpendicular to the optical plane were selected. Dotted lines indicate the apical surfaces, and arrowheads point to the apicalmost domain of the cell, which is magnified below. (G) Summary of the distribution of CAMSAP3 and MTs. CAMSAP3 is concentrated at the apicalmost region of the cell, associating with MTs. This MT population is reduced or absent in mutant cells. M, medial; D, dorsal; CTX, neocortex; V, ventricle; ChP, choroid plexus. Scale bars, 5 μm in (A), 2 μm in (C) and (F), and 1 μm in (D). See also Figure S6.

We then observed MT networks at the levels where N-cadherin-bearing AJs and BBs can be focused on. In WT ependymal cells with a monomeric centrosome, MTs radiated from it (Figure S6C). However, in cells undergoing centriole amplification, MTs not directly associated with BBs increased. In those with multiple BBs, MTs became organized in a meshwork over the BB and non-BB zones (Figure S6C). This MT meshwork was oriented in parallel to the apical cortex, unlike the perpendicular arrangement of MTs observed in intestinal epithelial cells (Toya et al., 2016). In CAMSAP3 mutant cells, MTs radiating from a centrosome or a small cluster of BBs were also observed. However, in those with multiple BBs, the MT meshwork tended to be less developed at the non-BB zone (Figure S6C).

Next, we closely analyzed MT organization in P7.5 WT ependymal cells along the apico-basal axis by serial optical sectioning of the images. At the apicalmost levels of WT ependymal cells, where CAMSAP3 was concentrated, MT filaments were detected, showing their association with CAMSAP3 puncta (Figures 6C-6E), as also confirmed by coronal sections of the cortex (Figure 6F). As expected, these MTs were located above the MT population associated with γ-tubulin^+^ basal bodies (Figure S6D). This apicalmost population of MTs was, however, depleted in CAMSP3 mutants (Figures 6D to 6F and S6D), although MTs were normally detectable below the CAMSAP3-localizing zones, where BBs are present. These observations suggest that CAMSAP3 produces apicalmost MT networks in ependymal cells and the lack of functional CAMSAP3 causes their depletion (Figure 6G).

## Discussion

Our present findings demonstrate that the CAMSAP3–non-centrosomal MT system is important for morphogenesis of the lateral ventricles. Loss of it brought about narrowing as well as increased stenosis or fusion of the ventricles. We sought primary causes of this ventricular deformation, finding that ependymal cells at the neocortex failed to undergo normal expansion of their apical domain, a final process of their differentiation. This failure apparently interfered with normal extension of the neocortical VSs along the mediolateral axis, and, in turn, widening of the ventricular cavity. These findings are consistent with previous observations that narrower ventricles are more prone to be stenosed under normal and pathological conditions (Ibañez-Tallon et al., 2004; Lowery and Sive, 2009; Shook et al., 2012).

CAMSAP3 proteins were concentrated at the apicalmost domain of ependymal cells, as observed in intestinal epithelial cells (Toya et al., 2016), suggesting that these different cell types share a common mechanism to place CAMSAP3 at particular subcellular sites. However, unlike in intestinal epithelial cells, the C-terminal domain-truncated CAMSAP3 also accumulated at similar sites in the case of ependymal cells, which suggests that non-functional CAMSAP3 behaves differently between these cells. At the subcellular regions where CAMSAP3 accumulated, MTs organized into networks that are oriented in parallel to the apical plasma membranes. In the absence of functional CAMSAP3, such MT networks were diminished. These observations suggest that CAMSAP3 serves to organize MT networks that are located at the apical-most regions of the cells and arranged in a horizontal orientation, although it remains to be determined whether it also regulates apico-basally oriented MTs as found in the intestines (Toya et al., 2016). On the other hand, BBs, which are another set of subcellular structures that associate with MTs, were located below the CAMSAP3-disributuing zone, and MTs present at the BB zones remained unchanged in CAMSAP3 mutant brains. A recent study showed that CAMSAP3 associates with BBs in a 1-to-1 fashion in multiciliated nasal epithelial cells (Robinson et al., 2020), but this kind of spatial organization of CAMSAP3 and BBs was not observed in ependymal cells.

We showed that mTORC1 signals became upregulated during ependymal differentiation and this process was important for their further growth. Our results suggest that the CAMSAP3-mediated MT networks likely support this process. It is known that mTORC1 activation occurs on lysosomes (Betz and Hall, 2013; Sancak et al., 2010), and, in CAMSAP mutants, lysosomes were less accumulated at the apical regions of ependymal cells, coinciding with the reduction of mTORC1 activity. Given that phospho-Akt, a kinase working upstream of mTORC1, was enriched in the apical portions of ependymal cells, these observations suggest that the lysosomal mispositioning in mutant cells might have been a cause of mTORC1 activity reduction. Thus, we propose a scenario that CAMSAP3 maintains a population of MTs which serves for redistribution of components required for mTORC1 signaling, such as lysosomes, and loss of this mechanism results in reduced mTORC1 activation, and in turn a failure of ependymal cell growth and the resultant narrowing of the lateral ventricle. One important question remains to be answered: why was VS growth less affected along the dorso-ventral axis of the brain? Alternative or additional mechanisms might be involved in VS extension at this side of ventricles, but cytological changes due to stenosis and fusion in the mutant ventricles prevented us from pursuing this question further.

Ventricle closure and resultant loss of ependymal cells were accompanied by depletion of adult NSCs at the striatal V-SVZ, confirming that the normal configuration of ependymal layers is important for maintaining NSCs. The depletion of NSCs was possibly due to loss of direct access to the CSF or of ependymal supports for their maintenance.

We unexpectedly discovered that apical accumulation of CAMSAP3 coincided with the onset of ependymal growth which requires activation of mTORC1. Ependymal cells likely have a developmental program for coordinating these independent cellular systems to work together, in order to attain their maturation. Elucidating the molecular basis of such a program would provide deeper insights into the mechanisms of how cells orchestrate multiple cellular machineries for their polarization, differentiation and growth.

## Materials and methods

### Mice

A CAMSAP3/Nezha mutant mouse line, *Camsap3^dc/dc^,* was reported previously (Toya et al., 2016). Briefly, the genomic sequence that encompasses the 14th and 17th exons was flanked by loxP sites, and deleted by crossing the floxed mice with the β-actin Cre transgenic ones. The resultant mice were simply designated as CAMSAP3 mutant mice in this work. The mutant mice were analyzed after heterozygous mice had been backcrossed at least for four generations to C57BL/6N. Noon of the day at which the vaginal plug was detected was designated as embryonic day (E) 0.5, and E19.5 was defined as postnatal day (P) 0.5. For all experiments, male or female mice were used. The experiments using mice were performed in accordance with protocol(s) approved by the Institutional Animal Care and Use Committee of RIKEN Kobe Branch.

### BrdU administration

5-Bromo-2′-deoxyuridine (BrdU, Sigma) was dissolved in drinking water at a final concentration of 0.8 mg/ml and given to P28.5 mice for 7 consecutive days. Mice were sacrificed at P49.5.

### Rapamycin treatment

Rapamycin (Merck Millipore) was dissolved in ethanol and diluted in water containing 5% (v/v) PEG400 (Nacalai Tesque) and 5% (v/v) Tween80 (Nacalai Tesque) at the final concentration of 40 μg/ml. P6 mice were intra-peritoneally injected with either rapamycin at 1 mg/kg body weight or vehicle only once a day for consecutive days.

### Fixation and sectioning

Embryos were dissected and perfused with 4% (w/v) paraformaldehyde (PFA) in phosphate-buffered saline (PBS; 0.1 M, pH 7.4) prewarmed at 37°C. Postnatal mice were anesthetized with sodium pentobarbital (100-200 mg/kg body weight; Abbott) and transcardially perfused with 1% (w/v) PFA in 37°C PBS, followed by 4% (w/v) PFA in 37°C PBS (Miller, 1981). For CAMSAP3, α-tubulin or tyrosinated tubulin immunostaining, PHM buffer (60 mM Pipes, 25 mM Hepes, 2 mM MgCl_2_) (Schliwa and Van Blerkom, 1981; Tanaka et al., 2012) was used instead of PBS. Brains were removed, postfixed in the same fixative for 2 hours at room temperature (RT) and subsequently overnight at 4°C, and cryoprotected by immersion in 10-30% (w/v) sucrose series in PBS. Fixed brains were embedded in OCT compound (Sakura Finetek), quickly frozen on isopentane cooled with liquid nitrogen and cut coronally at 20 μm using a cryostat (HM500M, MICROM). For en face imaging of the VS of the neocortex or the striatum, a single coronal slice was cut at the level of the anterior edge of the callosal commissure at 600 μm for E18.5 brains, 800 μm for P7.5 brains and 1 mm for P14.5, P28.5 and P49.5 brains using a microslicer (DTK-3000W, Dosaka EM). Usually, the posterior cut surface of the slice contained the anterior commissure. Then, a piece containing the V-SVZ of the neocortex or of the striatum was dissected from the slice, embedded in OCT compound, frozen on isopentane cooled with liquid nitrogen and cut in parallel to the VS at 10 μm using the cryostat. Alternatively, the dissected V-SVZ was used as a whole-mount.

### Immunofluorescence staining and microscopical imaging

Antigen retrieval was performed by incubating sections in a citrate buffer (10 mM citrate, 0.05% Tween-20, pH6.0) for 10 min at 98°C heated with a microwave (MI-77; Azumaya) or incubating sections in HistoVT one (Nacalai Tesque) for 20 min at 70°C. For BrdU detection, sections were incubated with 2M HCl for 30 min at RT. After two-time washes with PBS, sections were blocked in PBS with 0.3% (v/v) Triton X-100 (PBSX) containing 10% (v/v) normal horse serum (NHS) for 1 hour at RT followed by incubation with primary antibodies diluted in PBSX containing 5% NHS overnight at 4°C. Sections were washed two times with PBS and incubated for two hours at RT with corresponding secondary antibodies and subsequently incubated for 30 min at RT with DAPI (Sigma). Stained sections were washed two times with PBS and mounted with either Fluorsave (Chemicon) or TDE (hcbi.fas.harvard.edu). Whole-mount immunostaining was performed essentially in the same way as described above, except that no antigen retrieval was performed. Stacks of images were taken along the z-axis at optimal intervals using a confocal laser scanning microscope (LSM780; Zeiss) with a 20x/0.80 NA and a 40x/1.30 NA objective lenses or using Airyscan (LSM880; Zeiss) with a 63x/1.40 NA and a 100x/1.46 NA objective lenses. Acquired images were processed using Zen (Zeiss) and Photoshop CS5 (Adobe Systems).

### Antibodies for immunofluorescence

The rabbit antibody against CAMSAP3 was generated as described (Tanaka et al., 2012) and the rat antibody against N-cadherin MNCD2 was generated previously (Matsunami and Takeichi, 1995). The following primary antibodies were purchased: mouse anti-α-tubulin (Sigma, T9026), rabbit anti-β-catenin (Sigma, C2206), mouse anti-β-catenin (BD Transduction Laboratories, 610153), rat anti-BrdU (Bio-Rad, OBT0030), rabbit anti-cleaved Caspase-3 (Cell Signaling Technology, 9661), mouse anti-γ-tubulin (Sigma, T6557), rabbit anti-γ-tubulin (Sigma, T3559), rat anti-Lamp2 (Abcam, ab13524), rabbit anti-mTOR (Cell Signaling Technology, 2983), mouse anti-NeuN (Merck, MAB377), mouse anti-parvalbumin (Sigma, P3088), rabbit anti-phospho-Akt (Ser473) (Cell Signaling Technology, 4060), rabbit anti-phospho-MLC2 (Thr18/Ser19) (Cell Signaling Technology, 3674), rabbit anti-phospho-S6RP (Ser235/Ser236) (Cell Signaling Technology, 4858), rabbit anti-S100β (Dako, IS504), mouse anti-tyrosinated tubulin (Sigma, T9028) and rat anti-VCAM1 (BioLegend, 105701). The following secondary antibodies were used: CF-488, 568 or 647 conjugated donkey anti-mouse, rabbit or rat IgG (Biotium) and Alexa Fluor-488, 568 or 647 conjugated goat anti-mouse, rabbit or rat IgG (Invitrogen).

### Image analysis

To quantify the extent of ventricle fusion and stenosis in coronal sections, sections that contained three structures—the callosal commissure, the striatum and the septum—were selected at 200 μm intervals. Typically, three sections were selected for each mouse. The combined length of the stenosed and fused regions were measured along the VS using Metamorph (version 7.7, Molecular Devices) and divided by the length of the striatal VS in each section. The calculated values were averaged for each mouse.

The length of the neocortical VS was measured essentially in the same way as described above. To normalize the difference in the hemisphere size among brains, we calculated the proportion of the neocortical VS to the mediolateral length of the hemisphere, which was defined by the line that started at the midline of the callosal commissure, extended in parallel to the mediolateral axis of the hemisphere and ended at the pia matter of the neocortex.

To quantify the extent of ventricle fusion and stenosis on the striatal whole-mount, the striatal VS was classified into three regions based on the appearance of S100β signals: the area lacking S100 β signals was regarded as the fused region, the area with strong and uniform S100 β signals as the stenosed region, and the remaining S100β^+^ area as the conventional region that faces to the open ventricle (see also Results section). The surface area of each region was measured using Metamorph and its proportion to the entire VS was calculated.

To quantify the PV signal intensities in ependymal cells on the striatal whole-mount, boxed regions were put along a line that was drawn so that it crossed the boundaries between the stenosed and conventional areas. The average signal intensity in each boxed region was measured using Metamorph.

To count the number of VCAM1+ clusters on the striatal VS whole-mount, we excluded the one-third anterior part as well as one-third dorsal part of the striatal VS where weak VCAM1 signals could be detected in S100β^+^ ependymal cells. VCAM1 signals in the remaining part were binarized and the number of VCAM1^+^ profiles larger than 7 µm^2^ were counted using Metamorph. The number was divided by the area of the conventional region or of the conventional and stenosed regions.

To quantify the number of BrdU^+^ cells in the striatal V-SVZ, coronal sections were selected as described above. Typically, five or six sections were selected for each mouse. BrdU signals were binarized using Metamorph and BrdU^+^ profiles larger than 17 µm^2^, which was about a half size of the nucleus, were recognized as BrdU^+^ cells. These cells were manually counted in the V-SVZ, except for BrdU- and S100β-doubly positive cells, which are unlikely NSCs. The V-SVZ was defined as a high-cell density region along the striatal VS and its length was measured. The number of BrdU^+^ cells per 1 mm-length V-SVZ was calculated in each section. The calculated values were averaged for each mouse.

To calculate the density of BrdU^+^ cells in the GCL of the OB, coronal sections that were anterior to the accessory OB and contained the OB V-SVZ were selected at 200 μm intervals. Typically, three or four sections were selected for each mouse. The 400 μm-wide boxed region spanning the entire depth of the GCL was placed at the center of the line connecting the dorsal and ventral edges of the medial GCL using Metamorph. BrdU^+^ cells, which were defined as described above, were manually counted and NeuN^+^ cells were automatically counted using ITCN plugin (imagej.nih.gov/ij/plugins/itcn.html) of Image J. The number of BrdU^+^ cells was divided by the number of NeuN^+^ cells in each section. The same calculation was done for all selected sections and averaged for each mouse.

To estimate the total number of BrdU^+^ cells in the entire GCL in a single section, the area of the entire GCL was measured using Metamorph and divided by the area of the GCL in the boxed region (see above). The calculated value was multiplied by the number of BrdU^+^ cells in the boxed region. The estimation was done for all selected sections and summed for each mouse.

To measure the size of the telencephalon and the OB, the dorsal view of the brain was captured using a stereoscopic microscope (M420; Leica) attached with a CCD camera (DC500; Leica). The telencephalon and the OB were outlined and their areas were measured using Metamorph.

Apical domain size of a cell was measured using Metamorph by tracing cell boundaries visualized by N-cadherin or β-catenin immunofluorescence.

To count the number of cells on the neocortical VS along the mediolateral axis, the 100 μm-wide boxed region that spans the entire mediolateral length of the VS was placed at the center of the apicobasal length of the neocortical VS. The number of cells in the boxed region was manually counted.

Phospho-S6RP or phospho-Akt signal intensities were measured in cells undergoing centriole amplification, which were recognized based on the appearance of γ-tubulin signals. Ten consecutive optical planes taken along the z-axis (namely the apicobasal axis) at 0.16 μm-intervals were selected for each cell so that the middle plane was placed at the plane where N-cadherin^+^ junctions were the best in focus. Cells on the VS sharply steep to the optical plane were excluded from the analysis. The average signal intensity was measured in each optical plane using Metamorph and among the highest value was recorded for each cell. The background fluorescence was separately measured and subtracted.

To quantify the proportion of phospho-S6RP^+^ cells in each cell type (cells with monomeric BB, amplifying centrioles or multiple BBs) or the proportion of each cell type in phospho-S6RP^+^ cells, cells showing the average signal intensity of phospho-S6RP higher than 10 luminance (arbitrary unit in Metamorph) were recognized as phospho-S6RP^+^ cells.

To quantify the phospho-Akt distribution, six optical planes taken along the z-axis at 0.48 μm-intervals were selected for each cell so that the middle plane was placed at the plane where N-cadherin^+^ junctions were the best in focus. The average signal intensity was measured in each optical plane using Metamorph, which was divided by the intensity at the plane of N-cadherin^+^ junctions. The distribution was calculated in each cell and averaged for each genotype.

Lamp2 signal intensities were measured using Metamorph in three optical planes at 0.48 μm-intervals taken along the z-axis from the apicalmost of the cell. In the case of cells located on the VS steep to the optical plane, the inclination was normalized. In such cases, the apicalmost domain of the cell as well as the apical edge of β-catenin^+^ junctions extended across the consecutive optical planes at 0.16 μm-intervals, each of which contained part of them (see Figure 5G). To measure the Lamp2 intensity in the apicalmost domain of such case, the region surrounded by visible segments of the apical edge of β-catenin^+^ junctions and a line drawn to connect the segments’ ends was regarded as a part of the apicalmost domain in each plane. Lamp2 intensity at the parts of the apicalmost domain was separately measured and summed. The line(s) was kept unmoved in the following basal-side planes; the region surrounded by the line(s) and the lateral edge of β-catenin^+^ junctions and being 0.48 μm-beneath the apicalmost region was defined as the subapical region in each plane. The same method was applied to the regions 0.48 μm-beneath the subapical regions. The background fluorescence was eliminated before measurement using Zen. The measured intensities were divided by the total areas of the apical domain.

### Western blotting

P7.5 neocortical tissues were suspended in ice-cold lysis buffer (20 mM Tris-HCl pH 7.5, 1 mM EGTA, 1 mM EDTA, 50 mM NaCl, 1% (v/v) glycerol, 1% (v/v) Triton X-100) containing 0.25 mM NaF, 1 mM PMSF, 1 mM NaVO_3_ and a proteinase inhibitors cocktail (Roche) and lysed with a sonicator (UR-20P; TOMY). After centrifuge at 20,000 ×g at 4°C for 10 min, the lysate was added with 2-fold concentrated sample buffer (125 mM Tris-HCl pH 6.8, 20% (v/v) glycerol, 2% SDS, 0.006% Pyronin Y, 1% 2-mercaptoethanol), separated in a 5-20% (w/v) gradient SDS/PAGE gel (SuperSep Ace; FUJIFILM) and transferred to a PVDF membrane (Millipore). Blots were blocked with Tris-buffered saline (TBS) with 0.05% (v/v) Tween-20 (TBST) containing 5% (w/v) skim-milk for 1 hr at RT followed by incubation with primary antibodies diluted in TBST containing 3% bovine albumin overnight at 4°C. Blots were washed three times with TBST and incubated for 1 hour at RT with peroxidase-tagged secondary antibodies. Chemiluminescence was obtained using Novex ECL Chemiluminescent Substrate Reagent Kit (Invitrogen) and detected with a CCD imager (RAS-4000; GE healthcare). The following primary antibodies were used: rabbit anti-mTOR (Cell Signaling Technology, 2983), rabbit anti-phospho-mTOR (Ser2448) (Cell Signaling Technology, 5536) and mouse anti-GAPDH (Merck, G8795). Scanned images were quantified using Metamorph after monochrome inversion.

### Statistics

Data are presented as scatter plot with mean ± standard error of the mean except for phospho-Akt distribution, which were presented as mean ± standard deviation. Statistical significance was determined by the student t-test.

## Acknowledgements

We thank Yoko Inoue for technical support and the RIKEN Kobe light microscopy facility for imaging experiments.

## Author contributions

T.K. and M.T. designed the experiments. T.K. performed the experiments with the technical support by H.S. and M.K. T.K. and M.T. wrote the manuscript.

---

**Supplementary Figure 1.**
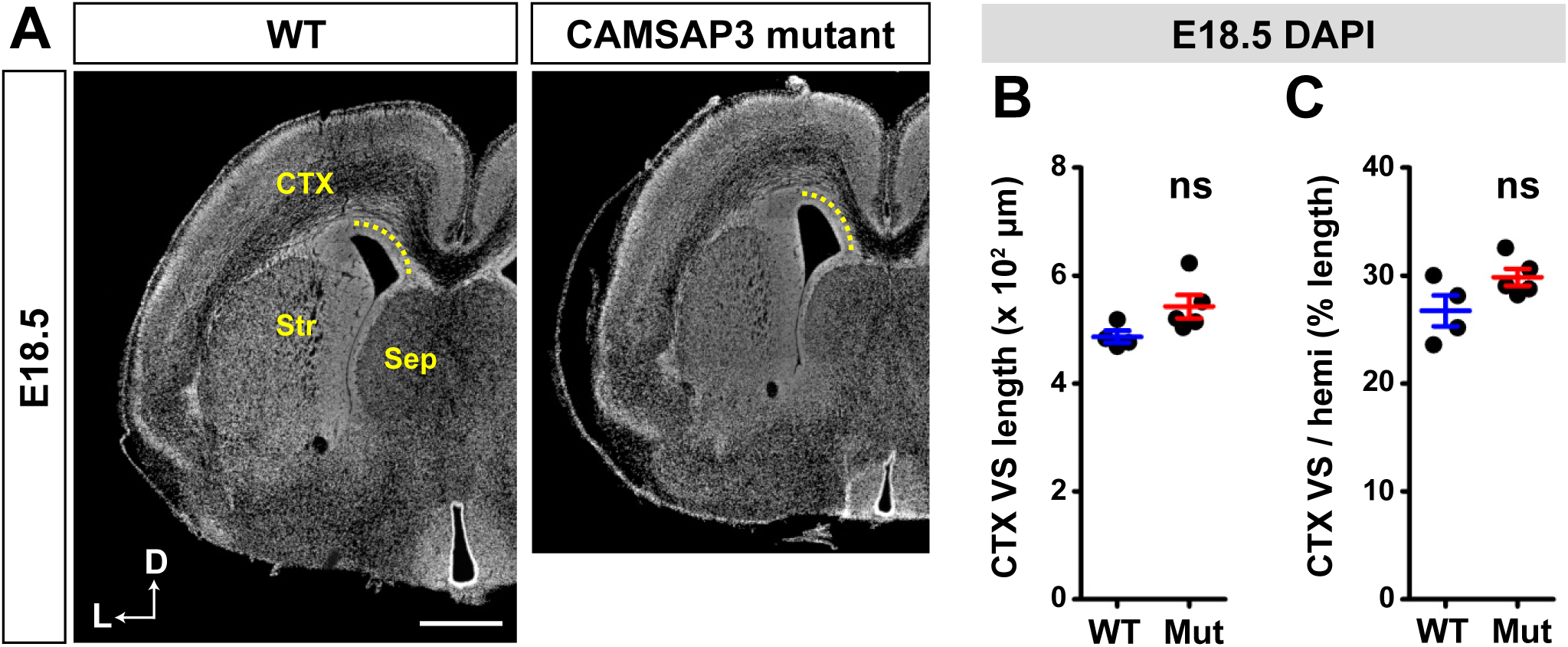
Embryonic brain of CAMSAP3 mutant mice. (**A**) Coronal sections of the WT or CAMSAP3 mutant telencephalon at E18.5, stained with DAPI. Dotted lines indicate the mediolateral width of the neocortical VS. (**B, C**) Mediolateral width of the neocortical VS (B), and the relative mediolateral width of the neocortical VS to the entire hemisphere (C) at E18.5. WT, n = 4; mutant, n = 5. D, dorsal; L, lateral; CTX, neocortex; Str, striatum; Sep, septum. Error bars indicate SEM. ns, not significant, Student’s t-test. Scale bar, 500 μm.

**Supplementary Figure 2.**
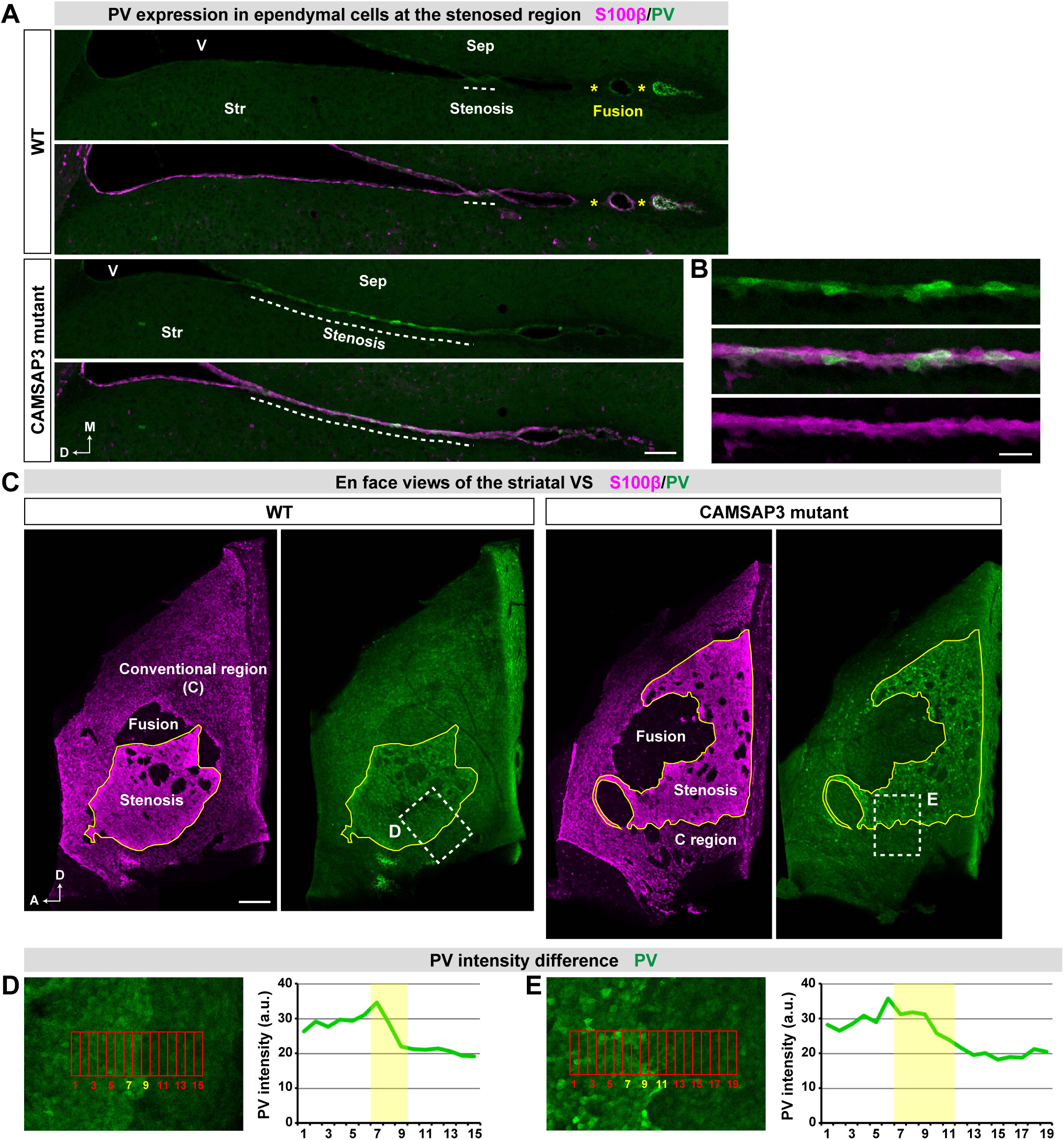
Parvalbumin distribution in ependymal cells at the stenosed region. (**A, B**) Coronal sections of the telencephalon at P28.5, immunostained for S100β and PV. Dotted lines and asterisks indicate the stenosed and the fused regions, respectively. In B, magnified views of ependymal cells at the stenosed region are shown. (**C**) En face view of a striatal VS whole-mounts at P28.5, immunostained for S100β and PV. The boxed regions are magnified in D and E. (**D, E**) PV immunostaining intensities at the stenosed and conventional regions. The average intensities in the boxes are plotted in the graphs where the boundary of the two regions is highlighted. M, medial; D, dorsal; A, anterior; V, ventricle; Str, striatum; Sep, septum. Scale bars, 100 μm in (A), 25 μm in (B), and 200 μm in (C).

**Supplementary Figure 3.**
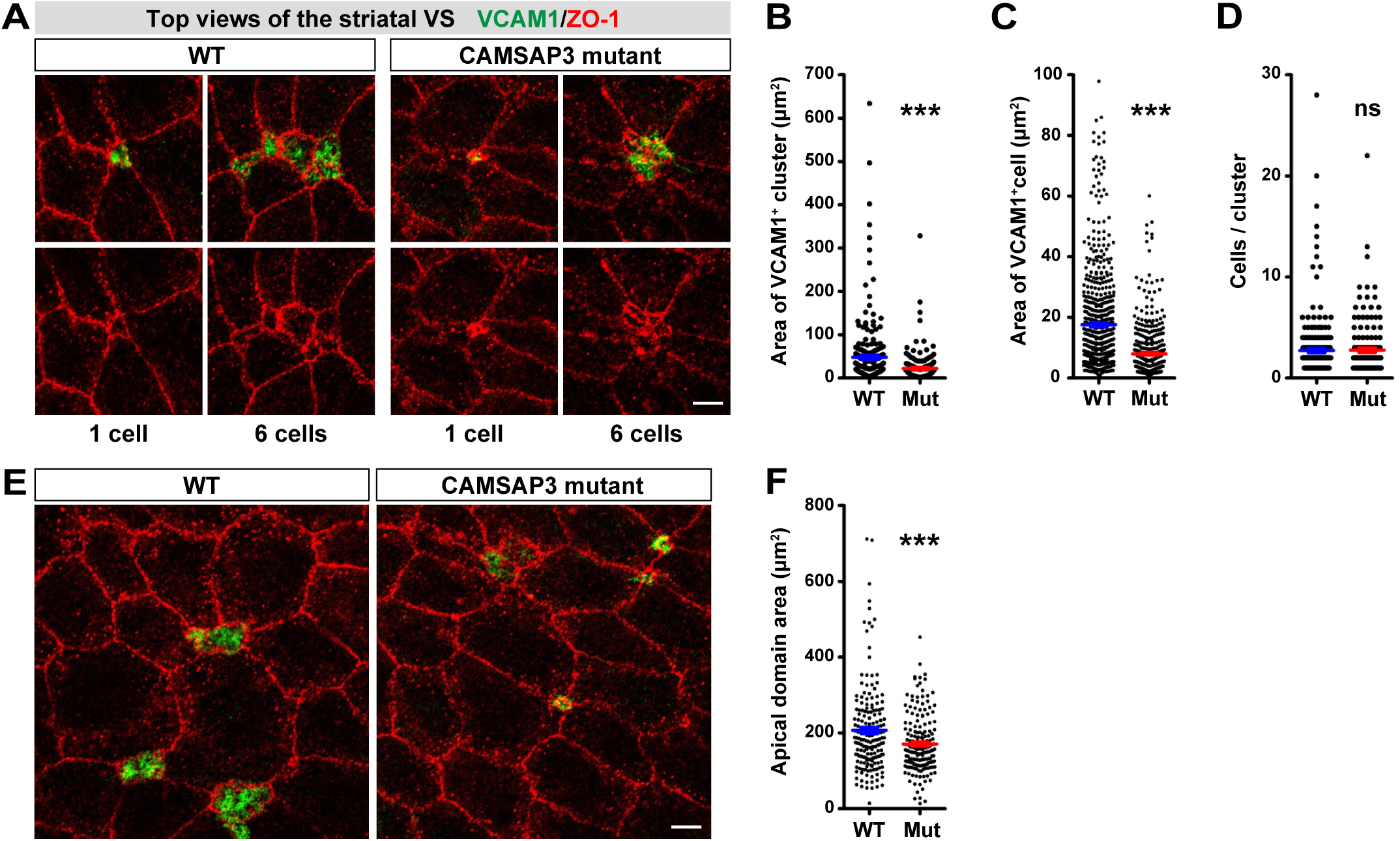
Apical domain areas of VCAM1^+^ cells and of ependymal cells at the striatum. (**A**) Top (en face) views of VCAM1^+^ cells in a striatal VS whole-mount at P28.5, immunostained for VCAM1 and ZO-1. Shown below are the numbers of cells per cluster. (**B**) Apical domain area of VCAM1^+^ clusters. n = 225 clusters in 2 brains for WT; n = 150 clusters in 2 brains for mutant. (**C**) Apical domain area of VCAM1^+^ cells. n = 614 cells in 2 brains for WT; n = 418 cells in 2 brains for mutant. (**D**) The number of cells per cluster. n = 225 clusters in 2 brains for WT; n = 150 clusters in 2 brains for mutant. (**E**) Top views of ependymal cells located in a dorsal part of the striatal VS whole-mount at P28.5, immunostained for VCAM1 and ZO-1. (**F**) Apical domain area of striatal ependymal cells. n = 60 cells per brain. 3 brains for each genotype. Error bars indicate SEM. ns, not significant, ***p < 0.001, Student’s t-test. Scale bars, 5 μm.

**Supplementary Figure 4.**
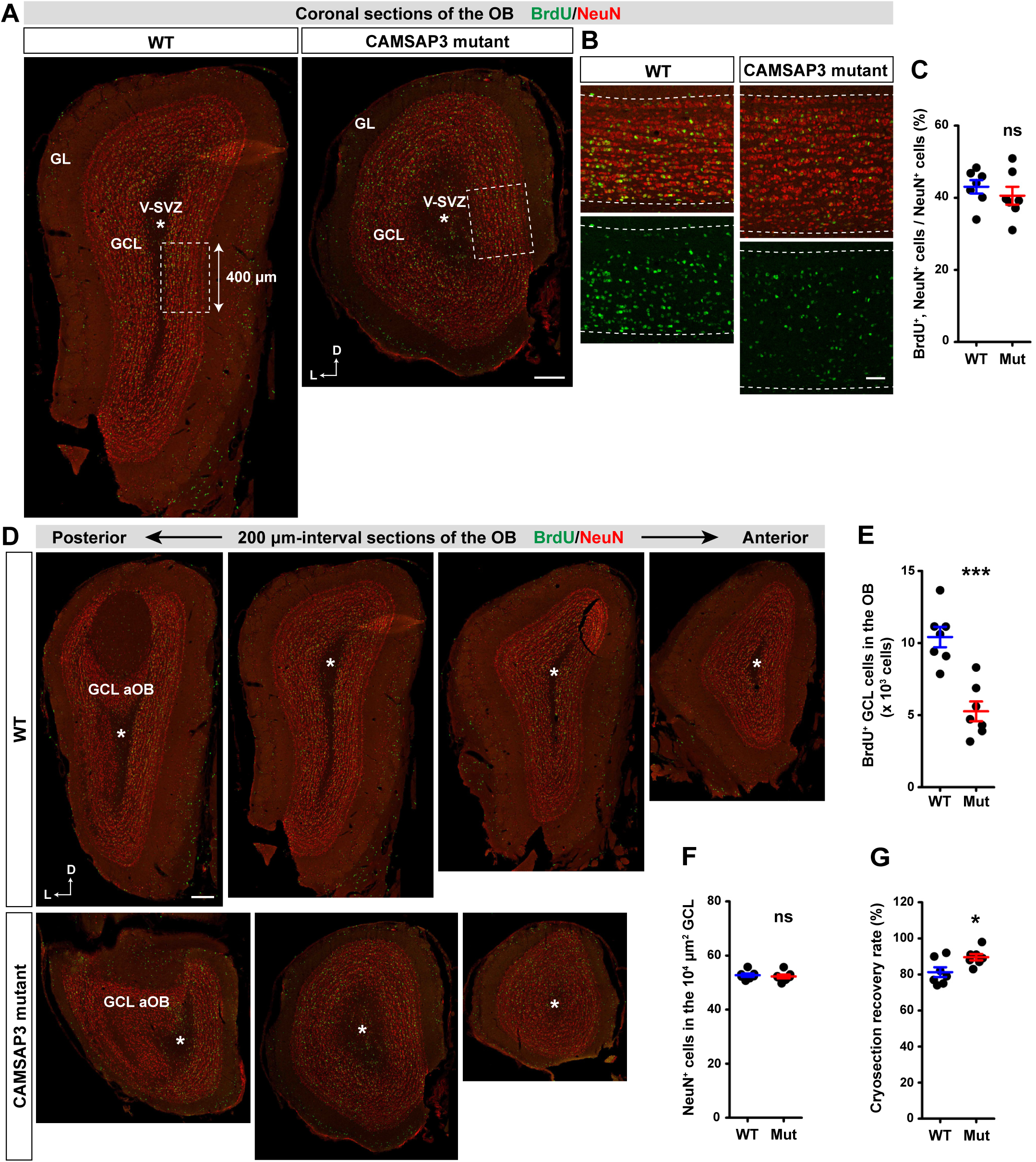
BrdU label-retaining cells in the OB at P49.5. (**A, B**) Coronal sections of the OB stained for BrdU and NeuN. Asterisks mark the V-SVZ in the OB. The boxed regions are magnified in B. Dotted lines indicate the boundaries of the GCL. (**C**) Percentage of BrdU^+^ cells in NeuN^+^ cells at the medial GCL. WT, n = 7; mutant, n = 7. (**D**) Consecutive coronal sections of the OB at 200 μm-intervals stained for BrdU and NeuN. Sections that are anterior to the accessory OB and that contain the V-SVZ were selected. Fewer sections were yielded due to the smaller OB in the mutant mice (see Figure 1A). Asterisks mark the V-SVZ. (**E**) The total number of BrdU^+^ cells in the entire GCL in the selected sections. WT, n = 7; mutant, n = 7. (**F**) Density of NeuN^+^ cells in the medial GCL. WT, n = 7; mutant, n = 7. (**G**) Recovery rate of cryosections. The fewer sections in the mutant OB are not a result of a lower recovery rate. WT, n = 7; mutant, n = 7. L, lateral; D, dorsal; GL, glomerular layer; GCL, granule cell layer; V-SVZ, ventricular-subventricular zone; aOB, accessory olfactory bulb. Error bars indicate SEM. ns, not significant, *p < 0.05, ***p < 0.001, Student’s t-test. Scale bars, 200 μm in (A) and (D), 50 μm in (B).

**Supplementary Figure 5.**
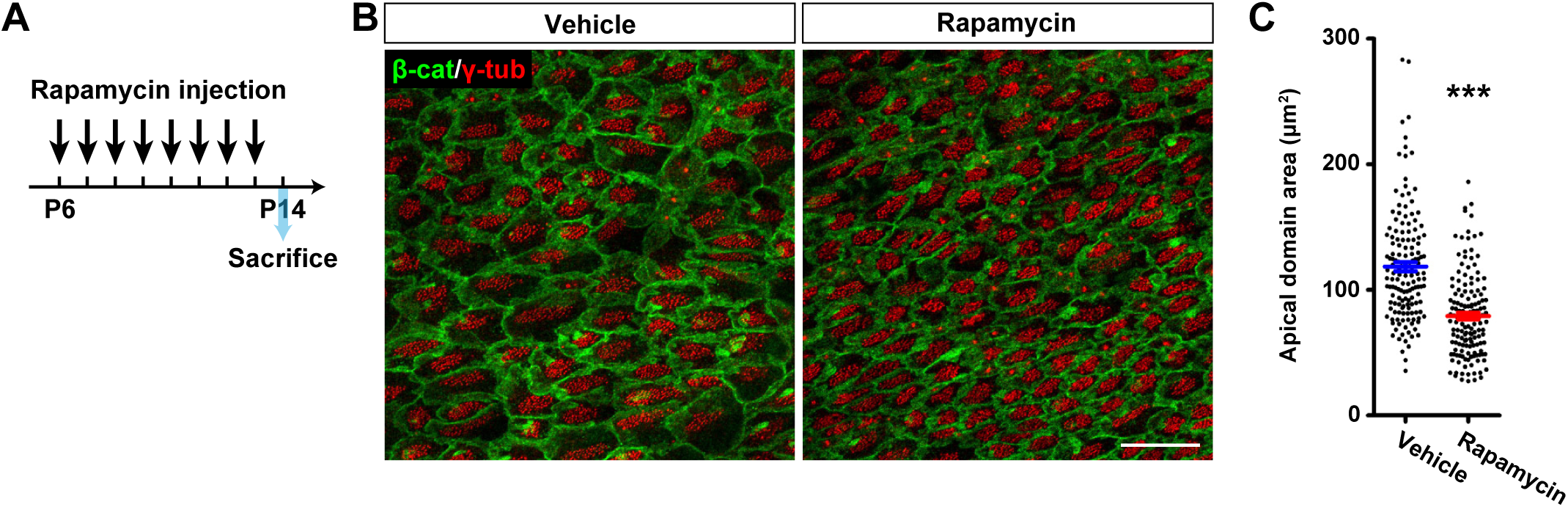
Inhibition of apical domain growth in neocortical ependymal cells by the mTORC1 inhibitor rapamycin. (**A**) Experimental timeline for rapamycin injections and sacrifice. (**B**) Top (en face) views of ependymal cells in a neocortical VS whole-mount of P14.5 WT mice that were injected with either the vehicle or rapamycin. Whole-mounts were stained for β-catenin and γ-tubulin. (**C**) Apical domain area of neocortical ependymal cells. n = 50 cells per brain. 3 brains for each condition. Error bars indicate SEM. ***p < 0.001, Student’s t-test. Scale bar, 20 μm.

**Supplementary Figure 6.**
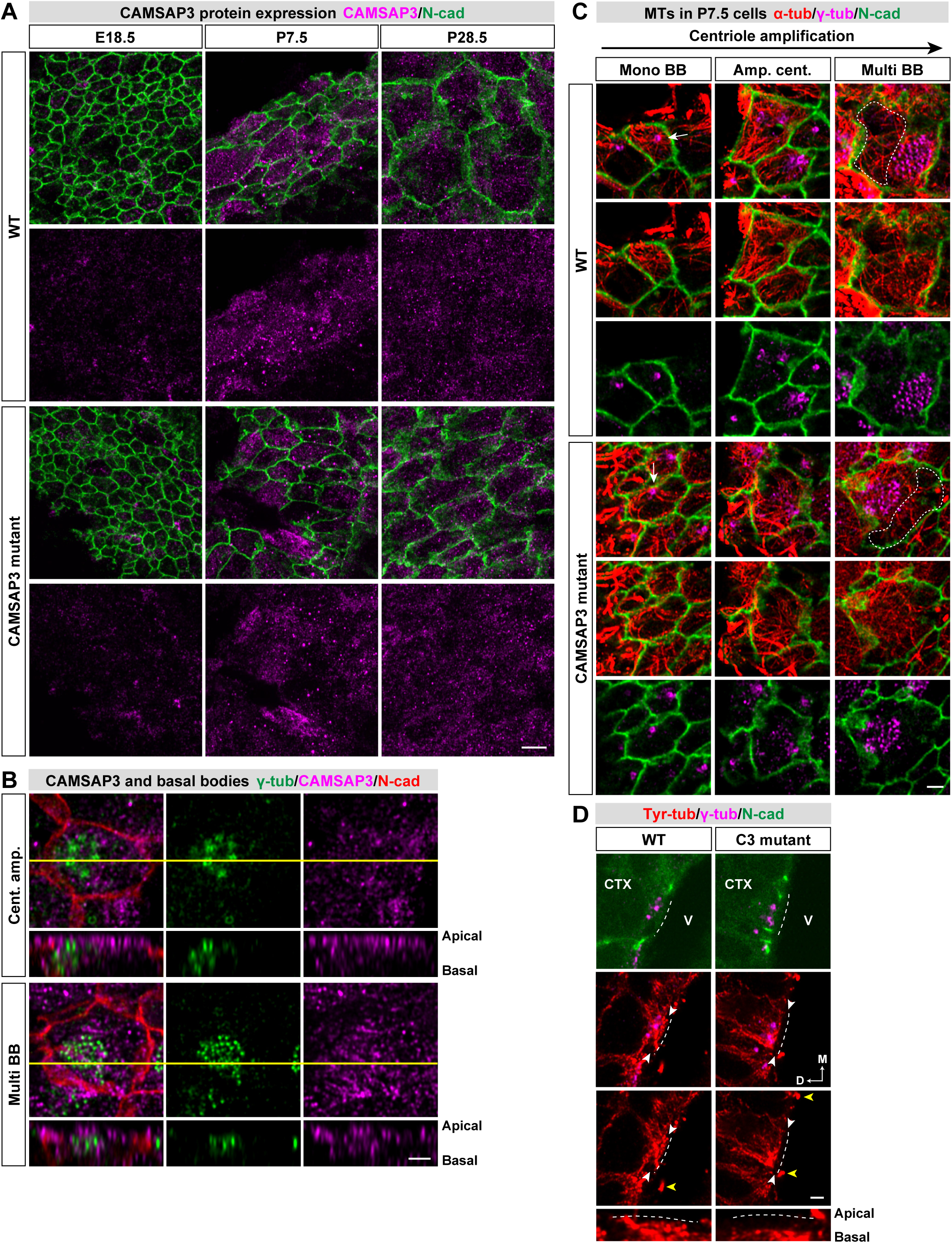
CAMSAP3 distribution and MTs in developing ependymal cells of the neocortex. (**A**) Top (en face) views of cells at the neocortical VSs at different ages, immunostained for CAMSAP3 and N-cadherin. (**B**) Top views of a WT cell with either amplifying centrioles or multiple BBs in the neocortical VS at P7.5, immunostained for γ-tubulin, CAMSAP3 and N-cadherin. The orthogonal (side) view at the line is shown below for each panel. (**C**) Top views of cells with monomeric BB, amplifying centrioles or multiple BBs in the neocortical VS at P7.5, immunostained for α-tubulin, γ-tubulin and N-cadherin. Single optical sections at the level where the apical edges of N-cadherin^+^ adherens junctions are focused on are shown. Non-BB regions are enclosed with broken lines. (**D**) Side (coronal) views of ependymal cells in the neocortical VS, immunostained for tyrosinated tubulin, γ-tubulin and N-cadherin. Cells undergoing centriole amplification were selected. The dotted lines indicate the apical surface, which is recognized by enhancing the background fluorescence in N-cadherin images. White arrowheads point to the apicalmost domain of the cell, which is magnified below. Yellow arrowheads point to cilia. Scale bars, 5 μm in (A), 2 μm in (B), (C) and (D).

## References

Baines, A. J., Bignone, P. A., King, M. D. A., Maggs, A. M., Bennett, P. M., Pinder, J. C. and Phillips, G. W. (2009). The CKK domain (DUF1781) binds microtubules and defines the CAMSAP/ssp4 family of animal proteins. Mol. Biol. Evol. 26, 2005–2014.

Banizs, B., Pike, M. M., Millican, C. L., Ferguson, W. B., Komlosi, P., Sheetz, J., Bell, P. D., Schwiebert, E. M. and Yoder, B. K. (2005). Dysfunctional cilia lead to altered ependyma and choroid plexus function, and result in the formation of hydrocephalus. Development 132, 5329–5339.

Betz, C. and Hall, M. N. (2013). Where is mTOR and what is it doing there? J. Cell Biol. 203, 563–574.

Bonifacino, J. S. and Neefjes, J. (2017). Moving and positioning the endolysosomal system. Curr. Opin. Cell Biol. 47, 1–8.

Delgado, A. C., Ferrón, S. R., Vicente, D., Porlan, E., Perez-Villalba, A., Trujillo, C. M., D’Ocón, P. and Fariñas, I. (2014). Endothelial NT-3 Delivered by Vasculature and CSF Promotes Quiescence of Subependymal Neural Stem Cells through Nitric Oxide Induction. Neuron 83, 572–585.

Doetsch, F., Caille, I., Lim, D. A., Garcia-Verdugo, J. M. and Alvarez-Buylla, A. (1999). Subventricular zone astrocytes are neural stem cells in the adult mammalian brain. Cell 97, 703–716.

Filice, F., Celio, M. R., Babalian, A., Blum, W. and Szabolcsi, V. (2017). Parvalbumin-expressing ependymal cells in rostral lateral ventricle wall adhesions contribute to aging-related ventricle stenosis in mice. J. Comp. Neurol. 525, 3266–3285.

Finlay, D. and Cantrell, D. A. (2011). Metabolism, migration and memory in cytotoxic T cells. Nat. Rev. Immunol. 11, 109–117.

Foerster, P., Daclin, M., Asm, S., Faucourt, M., Boletta, A., Genovesio, A. and Spassky, N. (2017). MTORC1 signaling and primary cilia are required for brain ventricle morphogenesis. Dev. 144, 201–210.

Furutachi, S., Miya, H., Watanabe, T., Kawai, H., Yamasaki, N., Harada, Y., Imayoshi, I., Nelson, M., Nakayama, K. I., Hirabayashi, Y., et al. (2015). Slowly dividing neural progenitors are an embryonic origin of adult neural stem cells. Nat. Neurosci. 18, 657–665.

Gao, Y., Moten, A. and Lin, H. K. (2014). Akt: A new activation mechanism. Cell Res. 24, 785–786.

Goodwin, S. S. and Vale, R. D. (2010). Patronin Regulates the Microtubule Network by Protecting Microtubule Minus Ends. Cell 143, 263–274.

Ibañez-Tallon, I., Pagenstecher, A., Fliegauf, M., Olbrich, H., Kispert, A., Ketelsen, U. P., North, A., Heintz, N. and Omran, H. (2004). Dysfunction of axonemal dynein heavy chain Mdnah5 inhibits ependymanl flow and reveals a novel mechanism for hydrocephalus formation. Hum. Mol. Genet. 13, 2133–2141.

Jiang, K., Hua, S., Mohan, R., Grigoriev, I., Yau, K. W., Liu, Q., Katrukha, E. A., Altelaar, A. F. M., Heck, A. J. R., Hoogenraad, C. C., et al. (2014). Microtubule Minus-End Stabilization by Polymerization-Driven CAMSAP Deposition. Dev. Cell 28, 295–309.

Kahle, K. T., Kulkarni, A. V., Limbrick, D. D. and Warf, B. C. (2016). Hydrocephalus in children. Lancet 387, 788–799.

Kokovay, E., Shen, Q. and Temple, S. (2008). The Incredible Elastic Brain: How Neural Stem Cells Expand Our Minds. Neuron 60, 420–429.

Kokovay, E., Wang, Y., Kusek, G., Wurster, R., Lederman, P., Lowry, N., Shen, Q. and Temple, S. (2012). VCAM1 is essential to maintain the structure of the SVZ niche and acts as an environmental sensor to regulate SVZ lineage progression. Cell Stem Cell 11, 220–230.

Korolchuk, V. I., Saiki, S., Lichtenberg, M., Siddiqi, F. H., Roberts, E. A., Imarisio, S., Jahreiss, L., Sarkar, S., Futter, M., Menzies, F. M., et al. (2011). Lysosomal positioning coordinates cellular nutrient responses. Nat. Cell Biol. 13, 453–462.

Kousi, M. and Katsanis, N. (2016). The Genetic Basis of Hydrocephalus. Annu. Rev. Neurosci. 39, 409–435.

Lasser, M., Tiber, J. and Lowery, L. A. (2018). The role of the microtubule cytoskeleton in neurodevelopmental disorders. Front. Cell. Neurosci. 12, 1–18.

Lim, D. and Alvarez-Buylla, A. (2016). The Adult Ventricular – Subventricular Zone and Olfactory bulb Neurogenesis. Cold Spring Harb. Perspect. Biol. 8, a018820.

Lowery, L. A. and Sive, H. (2009). Totally tubular: The mystery behind function and origin of the brain ventricular system. BioEssays 31, 446–458.

Martin, M., Veloso, A., Wu, J., Katrukha, E. A. and Akhmanova, A. (2018). Control of endothelial cell polarity and sprouting angiogenesis by noncentrosomal microtubules. Elife 7, 1–37.

Matsunami, H. and Takeichi, M. (1995). Fetal brain subdivisions defined by R- and E-cadherin expressions: Evidence for the role of cadherin activity in region-specific, cell-cell adhesion. Dev. Biol. 172, 466–478.

Meng, W., Mushika, Y., Ichii, T. and Takeichi, M. (2008). Anchorage of Microtubule Minus Ends to Adherens Junctions Regulates Epithelial Cell-Cell Contacts. Cell 135, 948–959.

Miller, M. (1981). Maturation of rat visual cortex. I. A quantitative study of Golgi-impregnated pyramidal neurons. J. Neurocytol. 10, 859–878.

Mirzadeh, Z., Merkle, F. T., Soriano-Navarro, M., Garcia-Verdugo, J. M. and Alvarez-Buylla, A. (2008). Neural Stem Cells Confer Unique Pinwheel Architecture to the Ventricular Surface in Neurogenic Regions of the Adult Brain. Cell Stem Cell 3, 265–278.

Muroyama, A., Terwilliger, M., Dong, B., Suh, H. and Lechler, T. (2018). Genetically induced microtubule disruption in the mouse intestine impairs intracellular organization and transport. Mol. Biol. Cell 29, 1533–1541.

Nashchekin, D., Fernandes, A. R. and St Johnston, D. (2016). Patronin/Shot Cortical Foci Assemble the Noncentrosomal Microtubule Array that Specifies the Drosophila Anterior-Posterior Axis. Dev. Cell 38, 61–72.

Paez-Gonzalez, P., Abdi, K., Luciano, D., Liu, Y., Soriano-Navarro, M., Rawlins, E., Bennett, V., Garcia-Verdugo, J. M. and Kuo, C. T. (2011). Ank3-Dependent SVZ Niche Assembly Is Required for the Continued Production of New Neurons. Neuron 71, 61–75.

Pongrakhananon, V., Saito, H., Hiver, S., Abe, T., Shioi, G., Meng, W. and Takeichi, M. (2018). CAMSAP3 maintains neuronal polarity through regulation of microtubule stability. Proc. Natl. Acad. Sci. U. S. A. 115, 9750–9755.

Pu, J., Guardia, C. M., Keren-Kaplan, T. and Bonifacino, J. S. (2016). Mechanisms and functions of lysosome positioning. J. Cell Sci. 129, 4329–4339.

Redmond, S. A., Figueres-Oñate, M., Obernier, K., Nascimento, M. A., Parraguez, J. I., López-Mascaraque, L., Fuentealba, L. C. and Alvarez-Buylla, A. (2019). Development of Ependymal and Postnatal Neural Stem Cells and Their Origin from a Common Embryonic Progenitor. Cell Rep. 27, 429–441.e3.

Robinson, A. M., Takahashi, S., Brotslaw, E. J., Ahmad, A., Ferrer, E., Procissi, D., Richter, C.-P., Cheatham, M. A., Mitchell, B. J. and Zheng, J. (2020). CAMSAP3 facilitates basal body polarity and the formation of the central pair of microtubules in motile cilia. Proc. Natl. Acad. Sci. 201907335.

Ruvinsky, I. and Meyuhas, O. (2006). Ribosomal protein S6 phosphorylation: from protein synthesis to cell size. Trends Biochem. Sci. 31, 342–348.

Sancak, Y., Bar-Peled, L., Zoncu, R., Markhard, A. L., Nada, S. and Sabatini, D. M. (2010). Ragulator-rag complex targets mTORC1 to the lysosomal surface and is necessary for its activation by amino acids. Cell 141, 290–303.

Sawamoto, K., Wichterle, H., Gonzalez-Perez, O., Cholfin, J. A., Yamada, M., Spassky, N., Murcia, N. S., Garcia-Verdugo, J. M., Marin, O., Rubenstein, J. L. R., et al. (2006). New neurons follow the flow of cerebrospinal fluid in the adult brain. Science 311, 629–632.

Schliwa, M. and Van Blerkom, J. (1981). Structural interaction of cytoskeletal components. J. Cell Biol. 90, 222–235.

Shook, B. A., Manz, D. H., Peters, J. J., Kang, S. and Conover, J. C. (2012). Spatiotemporal changes to the subventricular zone stem cell pool through aging. J. Neurosci. 32, 6947–6956.

Spassky, N., Merkle, F. T., Flames, N., Tramontin, A. D., García-Verdugo, J. M. and Alvarez-Buylla, A. (2005). Adult ependymal cells are postmitotic and are derived from radial glial cells during embryogenesis. J. Neurosci. 25, 10–18.

Takeichi, M. (2014). Dynamic contacts: Rearranging adherens junctions to drive epithelial remodelling. Nat. Rev. Mol. Cell Biol. 15, 397–410.

Tanaka, N., Meng, W., Nagae, S. and Takeichi, M. (2012). Nezha/CAMSAP3 and CAMSAP2 cooperate in epithelial-specific organization of noncentrosomal microtubules. Proc. Natl. Acad. Sci. U. S. A. 109, 20029–20034.

Toya, M., Kobayashi, S., Kawasaki, M., Shioi, G., Kaneko, M., Ishiuchi, T., Misaki, K., Meng, W. and Takeichi, M. (2016). CAMSAP3 orients the apical-to-basal polarity of microtubule arrays in epithelial cells. Proc. Natl. Acad. Sci. U. S. A. 113, 332–337.

Wu, J. and Akhmanova, A. (2017). Microtubule-Organizing Centers. Annu. Rev. Cell Dev. Biol. 51–75.

Yau, K. W., vanBeuningen, S. F. B., Cunha-Ferreira, I., Cloin, B. M. C., vanBattum, E. Y., Will, L., Schätzle, P., Tas, R. P., vanKrugten, J., Katrukha, E. A., et al. (2014). Microtubule minus-end binding protein CAMSAP2 controls axon specification and dendrite development. Neuron 82, 1058–1073.

